# What Markov state models can and cannot do: Correlation versus path-based observables in protein folding models

**DOI:** 10.1101/2020.11.09.374496

**Authors:** Ernesto Suárez, Rafal P. Wiewiora, Chris Wehmeyer, Frank Noé, John D. Chodera, Daniel M. Zuckerman

**Affiliations:** Advanced Biomedical Computational Science, Frederick National Laboratory for Cancer Research, Frederick, MD 21702; Computational and Systems Biology Program, Sloan Kettering Institute, Memorial Sloan Kettering Cancer Center, New York, NY 10065; Freie Universität Berlin, Germany; Department of Biomedical Engineering, Oregon Health and Science University, Portland, OR 97239

## Abstract

Markov state models (MSMs) have been widely applied to study the kinetics and pathways of protein conformational dynamics based on statistical analysis of molecular dynamics (MD) simulations. These MSMs coarse-grain both configuration space and time in ways that limit what kinds of observables they can reproduce with high fidelity over different spatial and temporal resolutions. Despite their popularity, there is still limited understanding of which biophysical observables can be computed from these MSMs in a robust and unbiased manner, and which suffer from the space-time coarse-graining intrinsic in the MSM model. Most theoretical arguments and practical validity tests for MSMs rely on long-time equilibrium kinetics, such as the slowest relaxation timescales and experimentally observable time-correlation functions. Here, we perform an extensive assessment of the ability of well-validated protein folding MSMs to accuractely reproduce path-based observable such as mean first-passage times (MFPTs) and transition path mechanisms compared to a direct trajectory analysis. We also assess a recently proposed class of history-augmented MSMs (haMSMs) that exploit additional information not accounted for in standard MSMs. We conclude with some practical guidance on the use of MSMs to study various problems in conformational dynamics of biomolecules. In brief, MSMs can accurately reproduce correlation functions slower than the lag time, but path-based observables can only be reliably reproduced if the lifetimes of states exceed the lag time, which is a much stricter requirement. Even in the presence of short-lived states, we find that haMSMs reproduce path-based observables more reliably.

## Introduction

The complexity of biomolecular stochastic dynamics presents significant challenges in extracting fundamental insight and building predictive models from atomistically-detailed molecular dynamics simulations. In the modern era of inexpensive graphics processing units (GPUs) and highly optimized molecular simulation codes capable of exploiting them, it is now routine to rapidly generate microsecond trajectories on a single GPU.^1–7^ Ready access to multiple GPUs now allows research laboratories to generate datasets tens to hundreds of microseconds in aggregate simulation time, ^8^ or with specialized supercomputers or distributed computing platforms, produce aggregate datasets over a millisecond in size.^9,10^ Distilling these enormous datasets into simple, mechanistic models capable of making predictions that can be confirmed experimentally and exploited for biophysical or pharmacological manipulation has been the focus of much of the field over the last decade.^11–13^

A particularly compelling approach has emerged in the machinery of *Markov state models*—discrete-state, discrete- or continuous-time stochastic models that approximate the stochastic dynamical evolution of biomolecules at equilibrium, coarse-grained in configuration space and time. ^11–19^ The essential ingredients of this model involve defining a set of conformational states representing regions of conformation space (*microstates*, defined in detail below) and a transition matrix that describes the probability of observing the system in a different state *j* some lag time *τ* after initially observing the system in state *i*. The availability of easy-to-use software tools for constructing Markov state models from molecular simulations^20^—especially PyEMMA^21,22^ and MSMBuilder^23–25^—have resulted in rapid uptake and widespread use of this technology;^11–13^ a Google Scholar search^1^ indicates over 500 papers were published referencing these models in 2018 alone, and over 3100 in total.

### Markov state models (MSMs) approximate the stochastic propagator of the biomolecular system

Despite their name, Markov state models (MSMs) do not assume the biomolecular dynamics must be truly Markovian once projected onto a discrete conformational state space—it is well-understood that the process of coarse-graining configuration space destroys the Markovian nature of the underlying stochastic dynamics in the full phase space of the system. Instead, MSMs aim to *approximate* the complex stochastic dynamics of the stochastic propagator or transfer operator of the system in a manner where the approximation error induced can be rigorously bounded by mathematical theory.^26–29^ Practically, constructing an MSM from a large quantity of simulation data requires a number of decisions to be made regarding choice of featurization of the molecular coordinates, selection of a dimensionality reduction scheme, and specification of a clustering strategy used to generate microstates; we refer to all of these choices as *hyperparameters* associated with MSM construction. ^12,13,18^ The complexity of hyperparameter selection has driven the development of software to automate the process of selecting appropriate hyperparameters from large combinatorial spaces,^30^ which necessitates the use of a numerical objective function to quantify model quality. By casting the problem of MSM construction in variational form, ^31,32^ the field has largely settled on the use of a quantity such as the variational approach to Markov processes (VAMP-*r*) score^32^ or generalized matrix Raleigh quotient (GMRQ)^33^ as an objective to be optimally maximized. To minimize statistical artifacts and penalize over-fitting, cross-validation is used to select optimal hyperparameters ^33,34^ subject to a fixed observation interval.^35^ Once optimal parameters have been selected, an appropriate lag time *τ* is selected using the timescales implied by the MSM constructed from different lag times (*implied timescales*, ITS).^18,36^ The model can then be used to describe statistical behavior, understand mechanisms, or predict properties on longer timescales than this lag time *τ*.^37^

### Markov state models induce approximation error in computed quantities

Coarse-graining of configuration space into discrete states introduces an approximation error into any property computed from the resulting MSM. ^26–29^ While this approximation error can be reduced either by increasing the number of (or optimizing the definitions) of the conformational microstates, the finite amount of trajectory data available usually means the primary means of reducing approximation error is to select a *lag time τ* large enough to incur minimal approximation error but small enough to ensure the model is capable of describing processes of interest that occur on timescales longer than *τ*.^18^ In the absence of statistical error, some computed properties will be exquisitely sensitive to this approximation error—for example, rate estimates may be highly sensitive to *τ*, and generally are too high^38^—while other properties, such as equilibrium properties, will be insensitive to it.

Despite the popularity of MSMs in extrapolating long-time dynamics from ensembles of short trajectories, there has not been a comprehensive assessment of the error (bias), sensitivity, and consistency in key observables estimated from MSMs as compared to computing these quantities directly from long MD simulations in protein systems. An analysis similar to ours was carried out for the Ala5 penta-peptide by Buchete and Hummer. ^16^ In particular, if an MSM is built from a very long MD trajectory, how do the MSM estimates of kinetic and path observables compare to those directly computed from the MD trajectory? The D. E. Shaw Research (DESRES) protein folding trajectories reported in Lindorff-Larsen et al.^39^ provide an opportunity for this comparison because several of the proteins exhibit > 10 transition events and hence accurate benchmark observables. Although MSMs have previously been built using the DESRES trajectories,^40,41^ those studies did not attempt the same type of quantitative analysis presented here using MSMs validated by modern techniques.

### History-augmented Markov state models attempt to resolve issues caused by coarse-graining

Recently proposed ‘non-Markovian’ or history-augmented MSM variants (haMSMs)^2^ attempt to overcome some of the challenges ordinary MSMs face in modeling certain properties of interest, ^44,45^ including mean first-passage times (MFPTs).^41–43^ The haMSMs build on prior work for history-tracing of trajectories^16,46^ and are similar to ‘exact milestoning’.^47^ These models can be built from the same trajectory data used to construct an ordinary MSM, but require specification of two (or more) macrostates of interest (e.g., folded and unfolded), based upon which kinetic observables will be computed; equilibrium properties of haMSMs, such as state populations, are identical to those of the corresponding MSMs. haMSMs condition the transition matrix on the macrostate that has been visited most recently (if at all) in any given trajectory, and this history information enables more accurate estimation of non-equilibrium observables related to macrostate transitions, particularly at short lag times when typical MSMs fail to exhibit truly Markovian behavior. For example, if every trajectory has visited at least one of the states and sampling is sufficient, a haMSM is guaranteed to yield the *exact* MFPT consistent with the time discretization of the raw trajectory data—regardless of the choice of microstates and using the shortest possible lag equivalent equivalent to the trajectory frame rate (i.e., setting *τ* = Δ*t*, see Table 1).

**Table 1:**
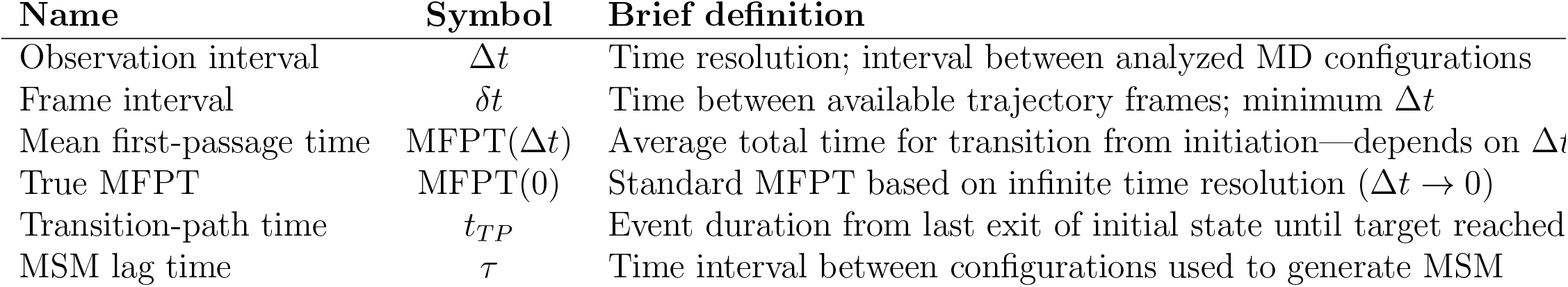
Key timescales pertinent to the present study. Each is defined more precisely in the main text, and some are depicted in Figure 1.

| Name | Symbol | Brief definition |
| --- | --- | --- |
| Observation interval | $\Delta t$ | Time resolution; interval between analyzed MD configurations |
| Frame interval | $\delta t$ | Time between available trajectory frames; minimum $\Delta t$ |
| Mean first-passage time | MFPT($\Delta t$) | Average total time for transition from initiation—depends on $\Delta t$ |
| True MFPT | MFPT(0) | Standard MFPT based on infinite time resolution ($\Delta t \rightarrow 0$) |
| Transition-path time | $t_{TP}$ | Event duration from last exit of initial state until target reached |
| MSM lag time | $\tau$ | Time interval between configurations used to generate MSM |

### The bias and accuracy of Markov state models can be assessed by comparison to long trajectories

Given recent developments in the field, this study attempts to fill a gap in the current literature via careful ‘apples-to-apples’ comparisons of MFPTs, rate constants, and dominant mechanisms from validated MSMs and history-augmented variants to the same observables calculated from long MD trajectories for protein (un)folding. We attempt to carefully control for several key factors: (a) the construction procedure for macrostates among which rates are computed, to avoid subjective choices as much as possible; (b) validation of the MSMs and the choice of the lag time—i.e., time discretization of the models; and (c) the formalism for estimating the rate, focusing on mean-first-passage-time and time correlation derived rates; (d) quantification of mechanism via a uniform approach for both MSMs and MD. For every observable, we attempt to account for the statistical power of the data through appropriate error bars.

Our use of carefully constructed macrostates enables quantification of *specific timescales* for transitions of interest, such as folding and unfolding, which contrasts with the more typical MSM-centric analysis of implied timescales (ITS) to identify slow processes that correspond to structural relaxation modes of the stochastic dynamics.^18,36,37^ Although ITS are mathematically well-motivated, the timescales identified this way may represent slow but uninteresting (improbable or spectroscopically silent) modes of the dynamics while the processes of interest may be much faster than the slowest identified timescales. For example, very slow partial unfolding events in a trajectory ensemble could mask a faster conformational exchange process of greater interest.

Our findings, on the one hand, confirm much of the promise of MSMs constructed from sufficient data using modern mathematical validation methods: at sufficient lag times, well-validated MSMs yield accurate kinetic predictions for the four proteins studied here. On the other hand, to achieve this fidelity, MSMs for the proteins considered here must utilize fairly long lag times *τ* ≳ 100 ns (see below and^48^) that coarse-grain temporal events faster than this timescale, which prevents the construction of credible mechanistic models of folding/unfolding pathways that could be compared head-to-head with mechanistic models derived directly from MD trajectories. As the transition time of folding events typically occur on much shorter timescales ≲ 10 ns, an MSM with a *τ* ≳ 100 ns lag time cannot reliably describe the statistics of these short events. Previous studies have noted the intrinsic limitation of MSMs for characterizing phenomena below the validated lag time.^18,36,49^ Another caution for future studies is the difficulty of obtaining a comparable quantity of trajectory data as was used in the present MSMs: smaller data sets could confound lag-time validation. Finally, when conformational states of interest can be defined as in the present systems, the haMSMs generally perform well even for short lag times and hence can provide accurate pathways as compared to MD for lag time matching MD time-discretization (*τ* = Δ*t*, Table 1).

## Theoretical Background

Although no new theoretical results or methods are presented in this report, here we briefly review essential background. Before describing the key elements of MSMs and haMSMs, we introduce general features of transition phenomena to assist readers in understanding the connections between the two approaches and the approximations employed.

We are concerned with a broad class of physical systems whose time evolution is described by trajectories **x**(*t*), where **x** denotes the set of all coordinates (such as a biomolecule and its solvent environment). Both equilibrium behavior (static and dynamic properties) and relaxation from out-of-equilibrium initial conditions could be estimated from a sufficiently large set of trajectories prepared in an appropriate way. We will consider systems of interest that evolve under stationary, thermostatted conditions, and obey detailed balance, such that a sufficiently long trajectory is guaranteed to reach equilibrium in a very long simulation.

### Microstates and macrostates

To construct a standard Markov state model (MSM), the whole of configuration space is first subdivided into a *partition of unity* ^3^, in which a a crisp division into regions called *microstates* is made. Each microstate is a compact, connected region of configuration space. We will follow MSM nomenclature in describing as a “microstate” a region small enough so that configurations within this region behave kinetically in a similar manner (and hence do not include large internal kinetic barriers between populated regions); the statistical dynamics should not strongly depend on which high-probability configuration within a microstate a trajectory is initiated from. Microstates are generally constructed from some sort of configurational clustering process of the sampled configurations—here, we use clustering approaches available in PyEMMA^22^ as described below. We note that the potential violation of these assumptions, especially at shorter lag times, ^41,44,45^ is a key motivation for defining haMSMs.

A “macrostate” is a larger region of configuration space expected to embody a kinetically metastable region, where transitions among microstates within a macrostate should be much more rapid than transitions among microstates in different macrostates.^49^ These macrostates may contain a substantial fraction the equilibrium probability (perhaps *p*^eq^ ≳ 0.1), though they may also represent kinetically metastable but low-population states of interest. For convenience, macrostates in the present study will always consist of collections of microstates; these are constructed using either a hierarchical kinetic clustering scheme described below, or derived by the eigenvectors of the MSM in a manner that captures kinetically related microstates.

We note that using dynamical models that obey microscopic detailed balance at the level of single configurations implies “coarse balance” at equilibrium—i.e., a lack of net flow between any pair of arbitrary regions in configuration space. ^52^ Hence, given rate constants (transition probabilities per unit time) *k_ij_* among micro- *or* macrostates *i* and *j*, we have

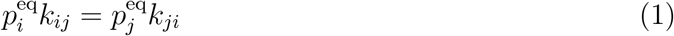

### Trajectories, transitions, and timescales

Trajectories **x**(*t*) may be usefully considered in different ways. A single long trajectory can be imagined which undergoes many transitions between arbitrary macrostates A and B, as shown in Figure 1(a). This trajectory can be decomposed into two directional components,^16,53,54^ the *α* subset consisting of segments currently or most recently in macrostate A, and the *β* component of segments currently or most recently in B. Roughly speaking, only the *α* components contribute to the dynamics/kinetics of the A → B transition and *β* to the B → A direction, although the two directions are necessarily related because of detailed balance.^55^

**Figure 1:**
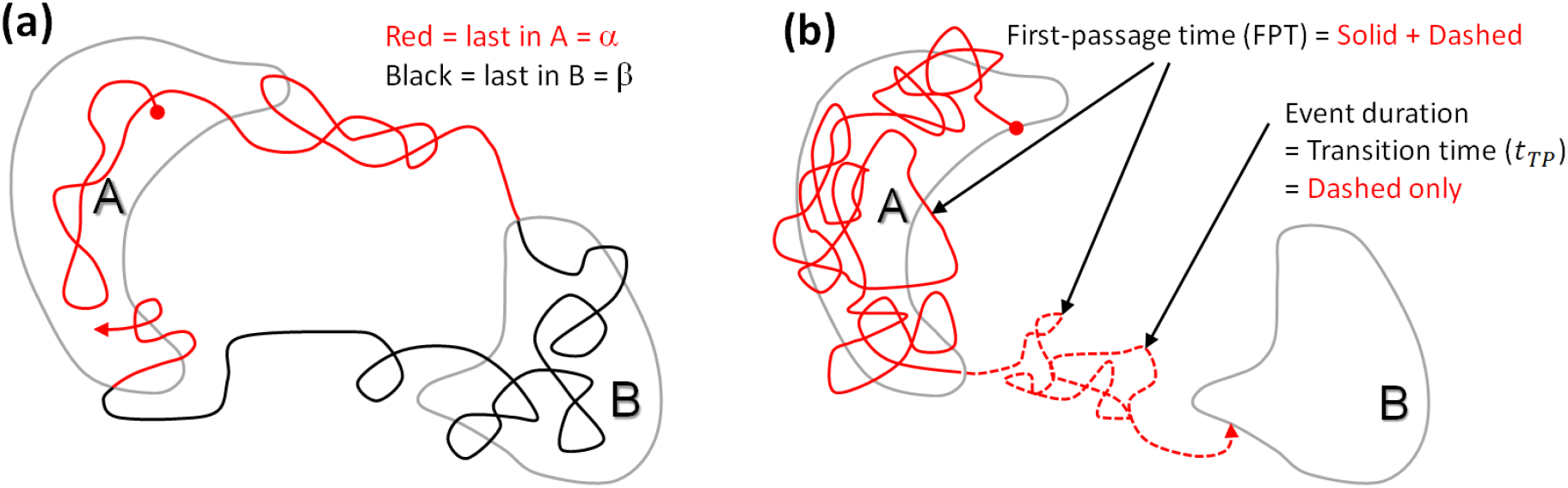
Illustration of trajectories, transitions, and first-passage times. Both panels are based on the arbitrary macrostates A and B (enclosed by grey lines) which are subsets of the configuration space represented by the plane of the page. **(a)** A single, very long trajectory exhibiting numerous transitions between macrostates can be decomposed into an *α* (red) component which contains all segments currently or most recently in macrostate A and a *β* (black) component, defined analogously for B. **(b)** A full A → B transition extracted from a long trajectory is characterized by the first-passage time (FPT), defined as the time elapsed from its first arrival to A (filled circle) until its first arrival at B (arrowhead). The segment of the trajectory following its last occupancy of A (dashed line) is known variously as the transition time, the event duration, the barrier-crossing time, or the transition-path time and will be denoted as *t_T P_*.

Below, we discuss the relevant classes of timescales of interest; all timescales and corresponding notation used in this study are briefly summarized in Table 1.

#### For metastable states, the mean first-passage time can be a useful way to characterize rates

Key timescales can be inferred by examining a segment of a long trajectory as in Figure 1(b). The first-passage time (FPT) for an A → B transition is defined as the elapsed time from when the trajectory first enters state A to when it first reaches state B, and analogously for B → A events. In practical situations, an FPT computed from a simulation trajectory necessarily will depend on the *observation interval* Δ*t*, the time between observed configurations: as Δ*t* increases, some first-entry events may be missed due to boundary recrossing and hence the FPT may monotonically increase; it cannot decrease.^41^ The average of all such FPTs in a given direction is the mean FPT (MFPT) for that direction, and in cases where states A and B define sufficiently metastable conformational states, the inverse MFPT quantifies a rate constant^52^ albeit one which generally is sensitive to macrostate definitions as seen below. Because the FPT depends on the observation interval Δ*t* so too will the MFPT—i.e., MFPT = MFPT(Δ*t*) and it will *monotonically increase* because of the missed events noted above.^41^

The traditional or mathematical MFPT corresponds to the MFPT(Δ*t* → 0) limit. All the trajectory data examined here is stored with a finite interval *δt* between ‘frames’ or configurational ‘snapshots’, so we will sometimes omit the argument from MFPT(Δ*t*) but readers should assume the Δ*t* dependence unless the Δ*t* → 0 limit is explicitly noted. In the case of the long trajectories considered here, snapshots were recorded with the interval Δ*t* =200 ps.

The MFPT is also expected to be sensitive to macrostate definitions in general. Consider the difference between describing a simple single-basin target state via a low or high iso-energy contour. Trajectories reaching a high-energy contour are much more likely to ‘bounce out’ of the state as compared to those reaching the low-energy contour. Stated more generally, some state definitions are less likely to suffer from re-crossing artifacts, but it must be borne in mind that for any given system, *there is no guarantee of the existence of physically well-defined states* characterized by fast intra-state dynamics and slow inter-state transitions. In the absence of such a separation of timescales, it should be noted that a system should not be characterized by a few-state kinetic model and estimating MFPTs may not provide physical insight.^56–58^

It is useful to understand the unphysical limit Δ*t* → ∞, or more practically Δ*t* » MFPT. In this scenario, frames are separated by a time interval longer than the (average) time for transitions, so the frames will appear to be a sequence of independent and identically distributed configurations, at least insofar as macrostate occupancy is concerned. Hence the probability of a configuration to occupy a given macrostate (A or B) is simply proportional to the equilibrium probability of the state at every time point, regardless of the previous configuration. Mathematically, we expect ^41^

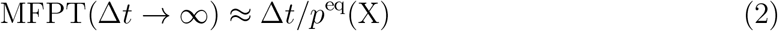

for transitions to state X = A or B characterized by equilibrium probability *p*^eq^(X)—i.e., simple linear behavior.

#### The transition-event time quantifies the duration of a rare transition

Another timescale of interest is the transition-event time (duration) *t_T P_* ^59–61^ which is defined to exclude the waiting time in the initial state: as shown in Figure 1(b), for the A→B direction, *t_T P_* is the duration of the final segment of a first-passage trajectory following the last visit to A, and analogously for the B → A direction.

Key timescales discussed in this study are briefly summarized in Table 1.

### Markov state model essentials

The overarching goal of Markov state modeling is twofold: First, describing the complex statistical dynamics of a stochastic biomolecular system with a simple discrete-state model that is both predictive of interesting properties, interpretable, and offers significant benefits for practitioners who might otherwise find themselves drowning in atomistic detail. Second, MSMs offer a way to bridge timescales by inferring model parameters from short trajectories that can then describe long-timescale behavior of the system, ideally offering a way around the need to directly simulate long-timescale or very rare events. When combined with adaptive sampling methods, ^62,63^ MSMs could in principle offer a highly efficient approach to the study of interesting slow biomolecular processes using only modest computational budgets. To achieve this, Markov state models aim to approximate the *transfer operator* of the underlying stochastic dynamics.^26–29^ The transfer operator 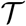 is defined in terms of its action on probability densities *p*(**x***, t*). In other words, if we prepared the system in some initial ensemble *p*(**x***, t*), waited a time Δ*t*, and then observed the ensemble *p*(**x***, t* + Δ*t*), what would the resulting ensemble look like? If we wait infinitely long, we reach the unique stationary distribution *p*^eq^(**x**) corresponding to thermodynamic equilibrium.

One way to constrain the complexity of a kinetic model is to construct *low-rank approximations* to the transfer operator. While the optimal fixed-rank approximation to the transfer operator is a linear combination of the eigenfunctions of the transfer operator, we do not know the eigenfunctions, and are forced to approximate them from the simulation data at hand. The Markov state model approach provides a principled way to construct these approximations, exploiting the metastability of the MD process. A metastable process corresponds to an approximately piecewise constant transfer function. Piecewise constant functions can be approximated by defining a partition on the configuration space into indicator functions (e.g., using clustering), and assigning each indicator function a weight. The standard MSM workflow ^11–13,17–19^ is to select a featurization, defining an appropriate distance metric (e.g., using tICA^40,64–66^), cluster snapshots to define microstates, and compute a transition probability matrix **T**(*τ*) between the resulting microstates, compute eigenvalues and eigenvectors of this matrix, and determine the earliest lag time *τ* at which it appears that the rate constants implied by this model are constant.

Once appropriately constructed, a key component of the Markov state model is the row-stochastic transition matrix **T**(*τ*), the row-stochastic matrix of transition probabilities among microstates:

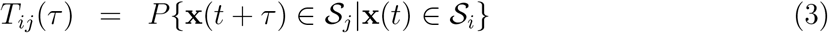

where 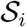 denotes the region of configuration space belonging to microstate *i*. These transition probabilities *T_ij_*, in contrast to rate constants *k_ij_*, refer to the probability that a trajectory which was in state *i* at time *t* will be in state *j* at time *t* + *τ*. As we assume the process is stationary, this transition probability *T_ij_*(*τ*) depends only on the lag time *τ* but not the origin observation time *t*. We further assume detailed balance with regard to the equilibrium probability 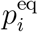, such that

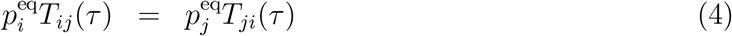

Once validated for a lag time *τ*, the time-dependent dynamics of the system can be estimated simply by exponentiating the transition matrix,

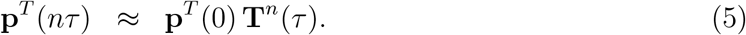

To improve the accuracy of such an approximation, we must either increase the resolution of our discretization in transition regions (increasing statistical noise or variance—as the transition probabilities between smaller, more rarely sampled partitions are harder to estimate) or increase the lag time *τ* (limiting our ability to resolve processes faster than the lag time).^26–29^

### Augmenting Markov models with history information

History-augmented MSMs (haMSMs) ^41^ avoid some of the assumptions of standard MSMs by separately making use of the *α* and *β* directional trajectory ensemble components (Figure 1) described above. The haMSMs employ history information which is always present in trajectories, but not used in standard MSM construction, when describing properties involving two conformational states of interest. Hence, a haMSM will exhibit different non-equilibrium properties compared to a MSM trained from the same data on the same microstates. Operationally, as detailed below, a haMSM is constructed by computing two sets of transition probabilities 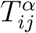 and 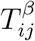 for the set of microstates by conditioning on *α* or *β* trajectory segments. Although the microstates of a haMSM need not be identical to those of a standard MSM for a given system, here we always construct MSMs and haMSMs using the same set of microstates.

The motivating idea for haMSMs can be seen more easily by employing a trajectory-ensemble perspective illustrated in Figure 2. An equilibrium ensemble of independent trajectories sufficiently long enough to connect states A and B can be decomposed into the directional *α* (last in A) and *β* (last in B) components.^16,53–55^ As the trajectory ensemble evolves in time, only the *α* component contributes to A→B transition behavior—timescales and mechanism—while only *β* generates B → A transitions. The “history” used in the haMSMs employed here is simply the *α* or *β* label, which potentially allows a more accurate description of a directional process because *β* trajectories do not participate in A → B transitions and can be excluded from their analysis, and likewise for the B → A direction. This concept is closely related to the *trajectory-based assignment* introduced by Buchete and Hummer^16^ and *core sets* used in transition interface sampling calculations, ^46^ which has been shown to permit a superior approximation of the slow eigenfunctions of the propagator due to its ability to implicitly approximate the committor functions between the states A and B.^26,29,50,51^

**Figure 2:**
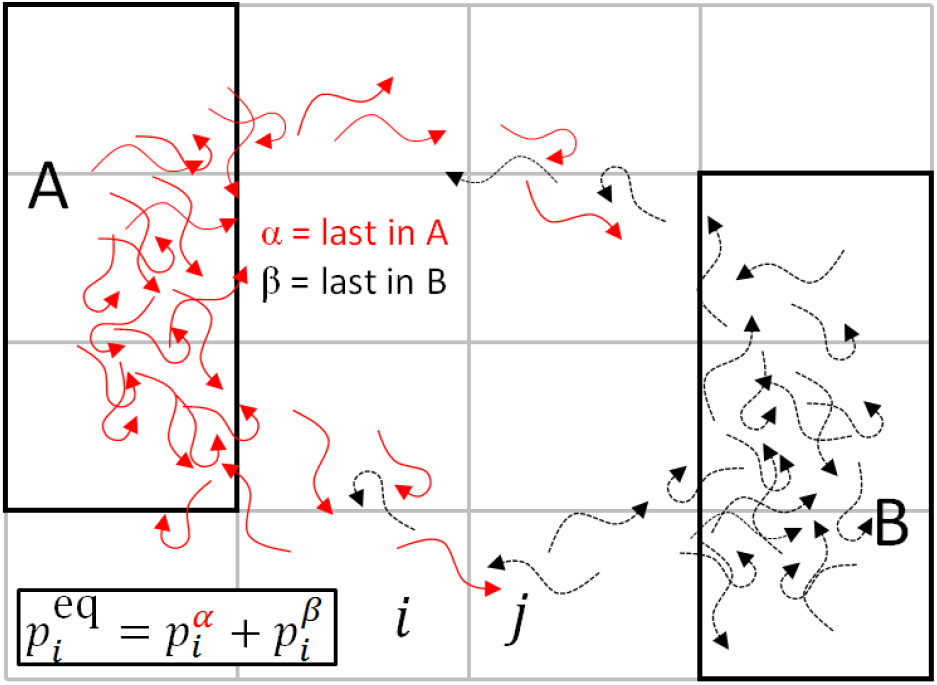
The equilibrium trajectory ensemble can decomposed into directional components that are used separately in an haMSM. The equilibrium ensemble of trajectories is a collection of independent and uncorrelated trajectories evolving for a very long time under the fixed conditions of interest. Each trajectory can be categorized according to the last-state scheme described in Figure 1: *α* (red) for those most recently in macrostate A and *β* (black) for those most recently in B. Hence every microstate (small rectangular cell) contains a mixture of both *α* and *β* trajectories. By definition, only *α* trajectories participate in A→B transitions; B → A transitions involve only *β* trajectories.

#### Equilibrium static quantities computed from a haMSM exactly match those of the corresponding MSM by construction

In principle, an arbitrary number of macrostates and corresponding trajectory types (*α, β, γ, …*) could be used to construct a haMSM, but more states will increase statistical noise given a fixed amount of data. In practice, users are more likely to investigate multiple two-state haMSMs to minimize statistical error.

The two-state *α/β* decomposition of trajectories in turn leads to decomposition of derived quantities such as equilibrium populations and rates. ^42^ In particular, for any microstate *i*, the equilibrium population is divided into the two directional components

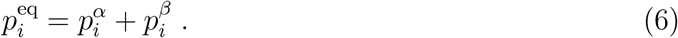

Note that because the equilibrium probabilities are normalized 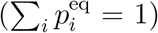, the sum over *α* or *β* populations separately are *not* normalized:

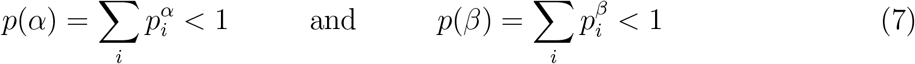

but they do comprise all trajectories so that *p*(*α*) + *p*(*β*) = 1.

In a similar decomposition, the transition probability *T_ij_* characterizing the overall transition rate in equilibrium (i.e., of a standard MSM) is also decomposed into a simple weighted average,

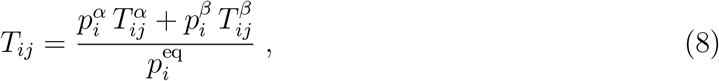

where 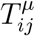 is the transition probability based only on the *μ* = *α* or *β* directional component. The relation (8) guarantees that static equilibrium properties derived from the set of *T_ij_*, such as state populations, will agree between a standard MSM and a haMSM using the same microstates.

The *α/β* decomposition is naturally related to the well-known committor analysis in a simple way.^67,68^ The committor, which is the splitting probability to reach a given state first (say, B) before another (A) starting from microstate *i*, is given exactly by 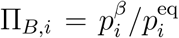. This follows from a reversibility argument: the next state to be reached is the time-inverse of the most recent state visited.^68^

For macrostate observables, the potential value of using the *α/β* decomposition can be readily understood for the MFPT. Based on the *exact* Hill relation^41,42,69,70^ between the MFPT and the (directional) steady-state probability flux into the target state, we have for the A→B direction

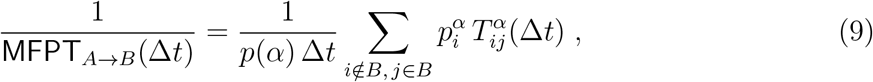

which is applicable whenever A and B are comprised exactly of sets of microstates. Importantly, the MFPT depends only on *α* properties and (9) is valid for *arbitrary* microstates regardless of whether they exhibit Markovian properties; likewise it is valid for arbitrary Δ*t*.^42^ In other words, the MFPT calculated from a haMSM trained on sufficient, unbiased trajectories will exactly match the MFPT which would be obtained from running a single long MD simulation and simply averaging FPT values, for any Δ*t* and arbitrary states. The *distribution* of FPT values generated from the haMSM is not guaranteed to match MD values, however. ^43^ Implicit in (9) is a requirement for consistency: to obtain the value MFPT(Δ*t*) from (9), the corresponding transition probabilities 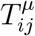 must be calculated using the same time discretization Δ*t*. For notational simplicity, the Δ*t* dependence of the MFPT and *T_ij_* will often be suppressed below.

In the realm of mechanism, the *α/β* decomposition in the haMSM again leads to exact *average* behavior, in terms of path fluxes. Specifically, the net flux from microstate *i* to *j* in the *α* component of the haMSM, 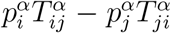, will exactly match the corresponding average *α* flux obtained from a very long MD simulation, and likewise for the *β* direction. Further, combining the *α* and *β* components of a haMSM will yield overall detailed balance, so long as detailed balance holds in the underlying equilibrium MSM. This can be seen by multiplying (8) by 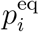 and comparing it to the corresponding index-reversed (*i* ↔ *j*) expression. However, haMSMs do not exhibit what might be called microscopic mechanistic reversibility. The ratio of probabilities of two individual *α* trajectories (defined as sequences of microstates) does not necessarily match the ratio for the reverse, *β* direction. In contrast, standard MSMs do exhibit microscopic mechanistic reversibility based on standard detailed-balance arguments.^71^

A haMSM can be used to compute any quantity available from a standard MSM. Some quantities can be calculated using analytic or recursive relations, such as the MFPT via (9). In general, arbitrary quantities defined on the space of discrete microstates can be computed for a haMSM using kinetic simulation based on the 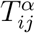 and 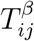 transition probabilities so long as the trajectory identity as *α* or *β* is tracked.

### Estimating transition probabilities in a haMSM

The history-labeled transition probabilities 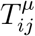 are a simple generalization of (3) and defined as

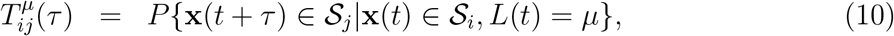

where the new element is the label operator, *L*(*t*) = *α* or *β*, which restricts consideration to one of the two trajectory subsets corresponding to the last macrostate visited. The estimation of 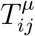 from unbiased MD trajectories typically is obtained from counting transitions

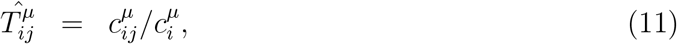

where 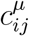 is the number of transitions observed (with label *μ*) from the microstate *i* to *j* at a given lag time *τ* and 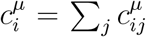. For convenience and in order to simplify the notation we are not showing explicitly the dependence of the transition probabilities and/or counts 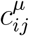 on *τ*.

In practical cases, only some transitions or trajectory segments can be traced back to a macrostate to yield the label *μ* = *α* or *β*. Trajectories insufficiently long to permit such labeling still can be fit into the haMSM formalism by generating a history label probabilistically, consistent with a Markov process. (Other strategies are possible, but we do not consider them here.) The likelihood of a label *μ* of a transition initiated at the microstate *i* is 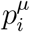 [recall (6)], and can be approximated through a Markov model in the absence of sufficient history. In concrete terms, if one imagines generating a very long discrete-state trajectory based on the Markovian matrix *T_ij_*, then for every *i* → *j* transition, the most recent macrostate can be traced back from the history of this trajectory; more simply, the 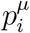 values could be computed directly from such a trajectory.

Thus, when traceback to a macrostate is not always possible, the haMSM transition probabilities are approximated by^72^

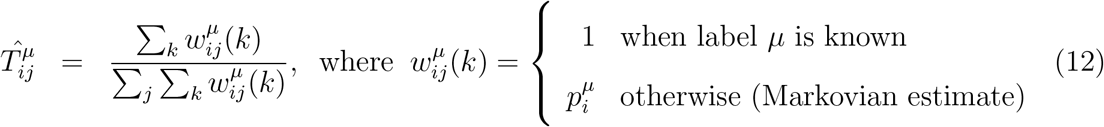

where the index *k* indicates summation over every instance of the *i* → *j* transition. In effect, each unlabeled transition is assigned fractionally to class *α* or *β* depending on 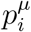. When all transitions are associated with a history label, (12) reduces to (11), as expected.

To obtain the 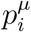 values analytically, the transition probabilities 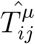 can be integrated in a single 2*N* × 2*N* row-stochastic matrix 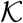, where *N* is the number of microstates. Then 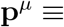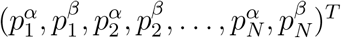 is the solution of 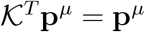.^41–43^ The Markovian approximation simply equates 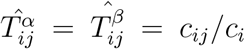, with the latter being unlabeled counts. In practice, we build 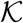 two times. First, we use the Markovian approximation as noted: although 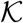 encodes the same model as **T** from (3) in this case, the Markovian estimation of 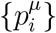 is straightforward from 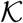. Then we obtain the final 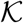—i.e., the haMSM–following (12).

### Related prior work

The analysis approach most closely related to haMSMs is the “core set MSM,” which in turn was motivated by the milestoning sampling strategy. ^73^ Core set MSMs were introduced by Buchete and Hummer^16^ and analyzed in mathematical detail by Schütte et al.^50^ Instead of requiring a full partition of the state space, core set MSMs require only some disjoint core-sets, ideally placed in the “cores” (kinetically central regions of high probability) of metastable sets of the dynamics. Trajectories are then “colored” according to which core they have most recently visited. In Ref.,^50^ the authors derive maximum likelihood and Bayesian estimators for the phenomenological rates (and for the finite-sampling error), present interpretations of the method in terms of Galerkin approximation, and note that the method does not require to choose a lag-time. They also note that the approximation quality of core set MSMs depends crucially on both the choice of core sets, and characteristics of the original dynamics. There has also been work on defining core sets automatically, using metastability-based,^74^ and density-based^75^ criteria, as well as further refinements to the concept of minimum dwell times required to constitute a core set visit. ^76^

Markov state models can also be estimated using a longer finite history. In,^77^ the authors propose a test for Markovianity at a particular lag-time by comparing the estimated transition probabilities of a first-order Markov process, *p*(**x**(*t*|**x**(*t* − *τ*)), with the predictions of second-order, *p*(**x**_*t*_|**x**(*t* − *τ*), **x**(*t* − 2*τ*)), or higher-order Markov processes, *p*(**x**_*t*_|**x**(*t* − *τ*), **x**(*t* − 2*τ*), …, **x**(*t* − *Nτ*)). Note that finite-order Markov processes can be reduced to first-order Markov processes on a suitably expanded state space.

The principle of retaining history information is also an implicit basis for a number of path-sampling approaches. Notably, the “transition interface sampling” method introduced the most-recent-state construction^46^ employed in haMSMs and in related work.^53,54^ The same concept is also embodied in the weighted ensemble method^42,68,78,79^ and forward flux sampling. ^80^

## Systems, macrostates, and MSM formulation

### Systems considered in this study

All analyses in this study were performed on the long equilibrium molecular dynamics trajectories of four miniproteins from the D. E. Shaw Research (DESRES) protein folding trajectories reported in Lindorff-Larsen et al.:^39^ chignolin, Trp-cage, NTL9, and villin (Figure 3). One trajectory was removed from the NTL9 dataset due to inconsistencies with the other three trajectories (see SI for details, which also discusses differing topologies within this dataset).

**Figure 3:**
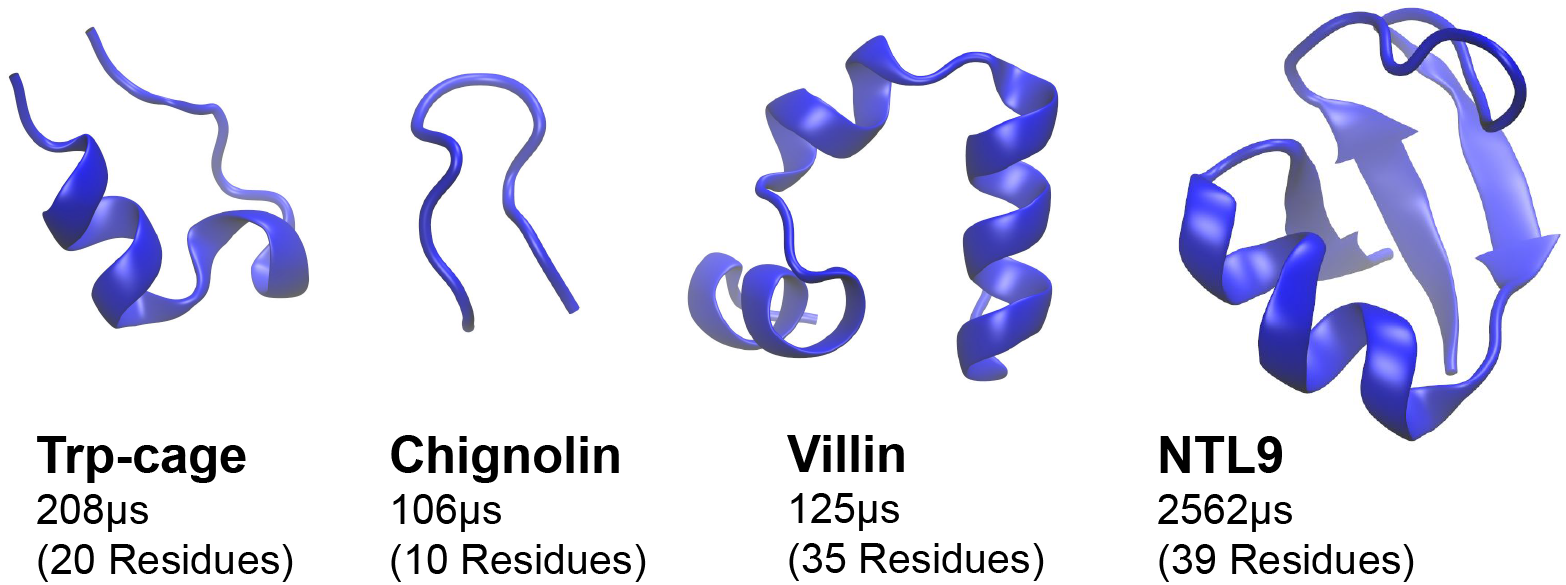
Protein systems considered in this study. All trajectory data comes from the D. E. Shaw Research (DESRES) protein folding trajectories reported in Lindorff-Larsen et al.^39^ The length of the MD simulation and the number of residues is specified in each case.

### MSM construction and validation

Numerous hyperparameters must be selected in the construction of Markov state models. ^30^ Key to the present study is the careful use of automatic hyperparameter selection selection ^30^ using an objective function derived from the variational approach to MSM construction ^31–33^ that uses cross-validation to ensure an optimal tradeoff between bias and variance. ^34^ While prior studies^40,41,81^ examined the fidelity with which MSMs reproduced the long-time behavior of some of these proteins, critically, they did not employ the most reliable validation procedures and hence the accuracy of those comparisons to MD studies may not be fully reliable.

#### Featurization and hyperparameter selection

To select the optimal MSM hyperparameters, we used variational scoring^31–33^ combined with cross-validation ^34^ to evaluate model quality, consistent with modern MSM construction practice. ^34^ To evaluate a large set of hyperparameters, reduced datasets subsampled to 10 ns/frame (50 ns/frame for NTL9) intervals were used for computational feasibility, except for chignolin, which remained at 0.2 ns/frame intervals due to its small size. The datasets were featurized with all minimal residue–residue distances (calculated as the closest distance between the heavy atoms of two residues separated in sequence by at least two neighboring residues). For consistency in interpretation and computational feasibility, this featurization choice was made without variational scoring. All parameters downstream of featurization (tICA lag time, number of tICs retained, tICA mapping, and the number of microstates) were then scored using a 100 ns MSM lag time; see Table 3 and the SI for further details and scoring results. We also explored using a much shorter MSM lag time of 10 ns, hypothesizing this could better optimize the reproduction of kinetics at short lag times, SI figures show the comparison of the results at the two scoring lag times.

**Table 3:**
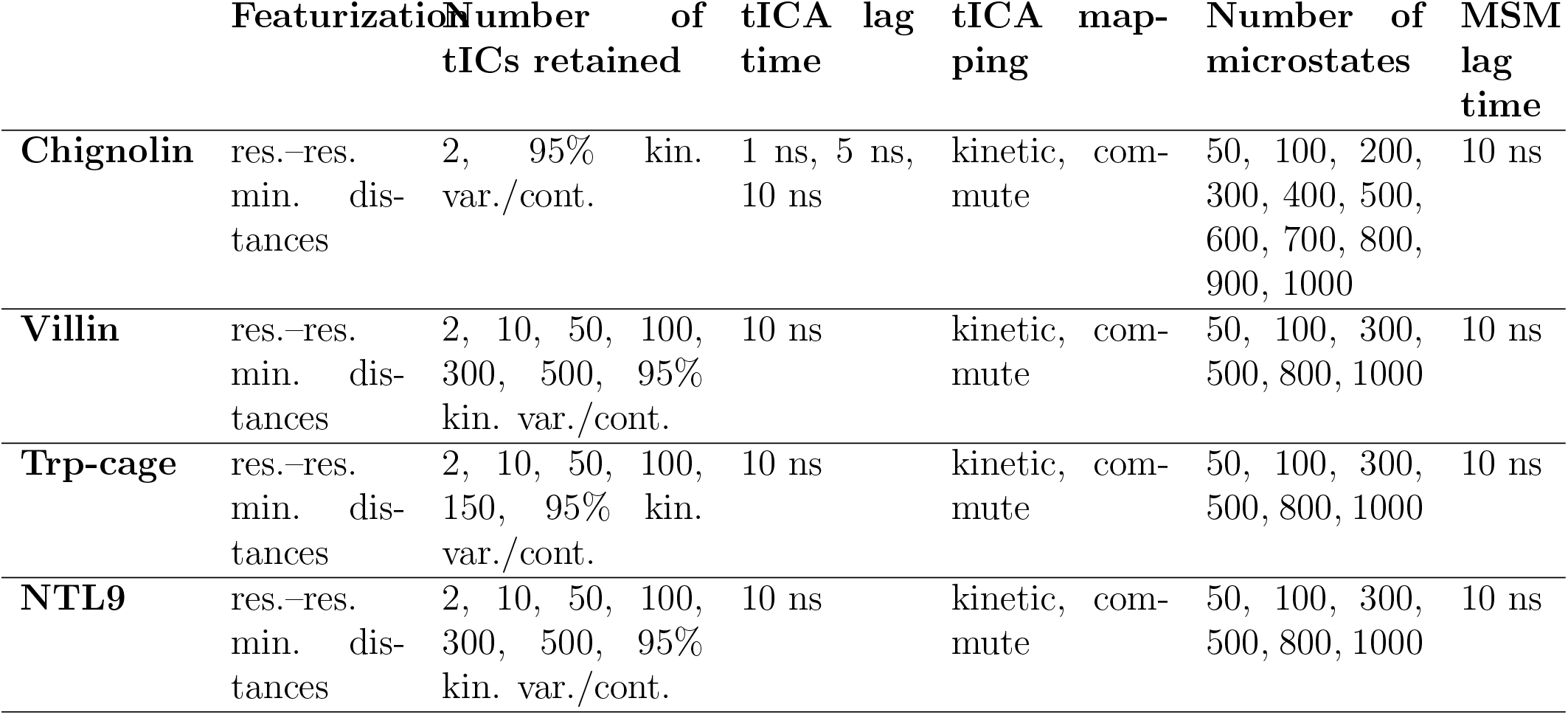
All of the model hyperparameters assessed combinatorially.

#### MSM scoring using cross-validation

We used a 50:50 shuffle-split cross-validation scheme to find the optimal set of hyperparameters while avoiding overfitting. In this scheme, 2 *μ*s long fragments of the trajectories (i.e., the original fragments in which the datasets are provided by DESRES) are randomly split into training and test sets of approximately equal sizes. tICA^40,64^ and k-means clustering were then conducted by fitting the model to the training set only, then transforming the test set according to this model. Scoring was based on the sum of squared-eigenvalues of the transition matrix (VAMP-2 score^32^), as this particular score is physically interpretable as ‘kinetic content’. Figure 12 shows the results of the scoring. Further details of the scoring procedure are discussed in the SI. To construct the discrete microstate trajectories used in this work, the modeling process was then repeated with full datasets (with no additional striding, i.e. at 0.2 ns/frame intervals and with no train-test splitting) using the top scoring parameters for the repeated tICA and k-means calculations.

#### Determination of useful MSM lag times

The convergence of the implied timescales in the final models was assessed by constructing Bayesian Markov state models (BMSMs) at increasing lag times (Figure 4). The following Markovian lag times at which the timescales first converged were identified: chignolin: 150 ns; villin: 100 ns; Trp-cage: 100 ns; NTL9: 200 ns. Chapman-Kolmogorov (CK) tests^18^ were conducted on the BMSMs to validate the self-consistency of the models at the Markovian lag times (Figure 14). CK tests were performed using two macrostates identified by PCCA++,^82^ except for villin where three macrostates were used (see SI for details).

**Figure 4:**
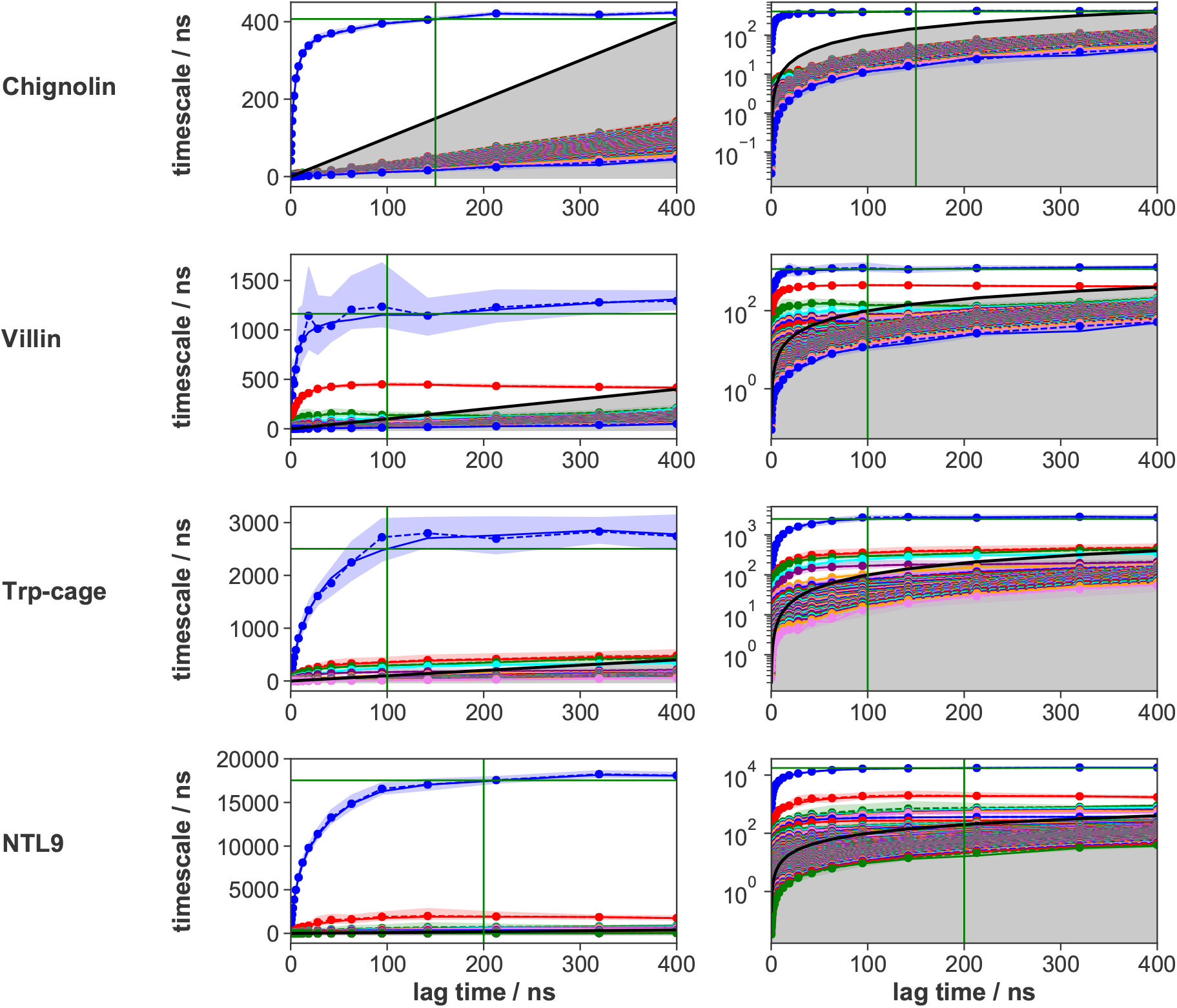
Implied timescales as a function of lag time for the Markov state models of all systems. All implied timescales of the BMSMs calculated at a range of lag times are shown: the maximum likelihood estimates (MLEs) as solid lines, the means of 100 samples as dashed lines, and the 95% confidence intervals of the means as shaded regions, estimated using Bayesian MSM methods implemented in PyEMMA. Note that the left panels use linear scale for implied timescales, while the right panels use logarithmic scale. The gray area signifies the region where timescales become equal to or smaller than the lag time and can no longer be resolved. Vertical green lines mark the lag times chosen for the MSMs used here, at which the timescales converge (chignolin: 150 ns, villin: 100 ns, Trp-cage: 100 ns, NTL9: 200 ns), while horizontal green lines mark the MLEs of the slowest timescales of MSMs computed at those lag times.

### Macrostate construction and the transition path time

We employed two different macrostate construction schemes, based on different clustering approaches, to ensure that the results of our study are not sensitive to the chosen process.

As our primary method, the MSMs were coarse-grained into two (“folded” and “unfolded”) macrostates, except for villin for which three macrostates (“folded”, “unfolded” and “misfolded”—see SI for details) were necessary, using the fuzzy spectral clustering method PCCA++.^82^ The identities of the macrostates were assigned based on visual inspection of chosen segments of the trajectories in PyMOL.^83^ The coarse-graining resulted in macrostates with the following equilibrium populations: chignolin: 79.4% folded, 20.6% unfolded; villin: 32.5% folded, 60.9% unfolded, 6.6% misfolded; Trp-cage: 19.0% folded, 81.0% unfolded; NTL9: 91.5% folded, 8.5% unfolded.

In order to study the mechanisms of folding, we sought to define an intermediate region, leaving more core-like folded and unfolded states. As PCCA++ produces fuzzy metastable membership of microstates into macrostates, we defined the intermediate region to consist of the 10% of all microstates with macrostate memberships closest to 50%, with the remaining 90% of microstates assigned to the folded/unfolded/misfolded macrostate to which they had the highest membership.

In the second macrostate construction procedure, we employed a hierarchical kinetic clustering procedure—a variant of a published process^84^ which in turn is based on an earlier proposal.^49^ Specifically, the clustering procedure is based on the commute time *t_ij_* between every pair (*i, j*) of microstates at the highest time resolution, i.e., *t_ij_* = MFPT_*i→j*_(*δt*) + MFPT_*i→j*_(*δt*), where *δt* is the time interval between available trajectory frames (minimum possible lag-time). Since direct estimation of *t_ij_* from MD data would be very noisy, we use Markovian MFPTs computed at that short lag-time. Our goal here is not to build the best model for computing *t_ij_* but a quick recipe for the construction of the macrostates. See SI for further details of the procedure.

### Software and code availability

PyEMMA 2.5.4 and 2.5.6^22^ was used for all MSM calculations. All code used for this analysis is available via a Github repository at https://github.com/choderalab/msm-mfpt.

## Results

### Transition-path time

The transition-path time *t_T P_* (Figure 1 and Section) provides a critical filter for understanding the domain of applicability of MSMs. Once macrostates have been defined, we can evaluate the transition-path time *t_T P_* —i.e., the duration of a transition event, from its last departure from the initial (e.g., unfolded) macrostate until its first arrival to the target (e.g., folded) state. Table 2 shows transition path times from MD simulation for the proteins typically are less than 10 ns, with NTL9 about an order of magnitude longer. The table values combine both folding and unfolding events because of the microscopic reversibility of MD trajectories.

**Table 2:** Transition event durations *t_T P_* (ns) from long MD simulations.

|  | chignolin | Trp-cage | villin | NTL9 |
| --- | --- | --- | --- | --- |
| median | 0.6 | 0.4 | 0.4 | 21.2 |
| average | 1.6 | 8.5 | 2.7 | 108.7 |
| std. dev. | 3.2 | 32.3 | 7.7 | 256.0 |

Because the validated MSM lag times generally are longer than the *t_T P_* values, the MSMs should not be used to probe intra-transition characteristics. That is, the validated lag time represents the finest time resolution for which the MSM can address relevant questions.

### Observable: MFPT analysis

The MFPT (Section) is a key characteristic of chemical and physical processes.^69,85,86^ Although sensitive to both lag time and macrostate definitions as noted above, the MFPT does quantify a well-defined physical process by construction, in contrast to implied timescales. This concreteness makes the MFPT an ideal yardstick for comparison among model and reference data.

Figure 5 shows the comparison of both MSM and haMSM predictions for the MFPT based on PCCA++ macrostates,^82^ as compared to reference MD results. Recall that the MFPT intrinsically depends on the time resolution Δ*t* which is taken to match the MSM lag time *τ*. All the proteins exhibit similar behavior. At short times below the validated MSM lag values, the MFPT shows strong lag-time sensitivity and MSMs are “faster” than MD; the haMSMs successfully track the MD behavior even in this regime. In the quasi-plateau region following the first inflection all the models become consistent with MD. At large times, after the second inflection, we see the trivial (non-kinetic) asymptotic linear behavior of (2). The haMSMs track the MD data in all the regimes, even when limited history information (Section) is used.

**Figure 5:**
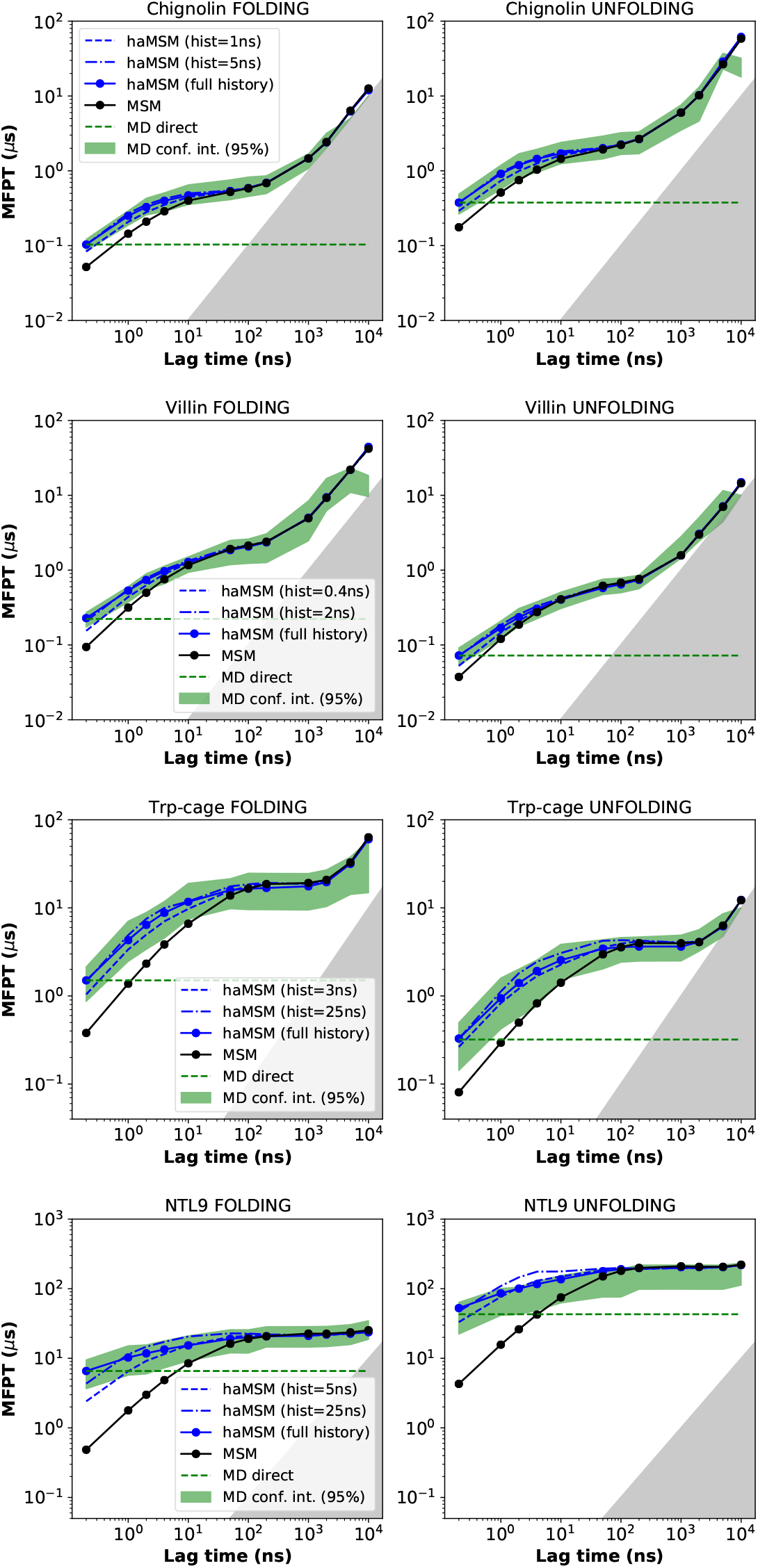
MFPT estimates compared among MD, MSMs, and haMSMs. The MFPT for both folding and unfolding is plotted as a function of lag time. Reference MD data is shown as the 95% confidence interval (green band), which can be compared to validated MSM data (black lines) and haMSM values with full history (solid blue lines) and partial history (dashed blue lines). The gray area signifies the region where MFPTs become equal to or smaller than the lag time and can no longer be resolved. The MD confidence intervals missing for the final data points of chignolin and villin are due to no more transition events seen at those very long lag times.

Comparing MSM predictions to MD behavior further validates the relatively long lag times required for these systems. In every case, the MSMs do not track the MD data until lag times *τ* ≳ 100 ns. We note that even for the MD data, the MFPT does not always reach a true plateau region where it is insensitive to lag time, which reflects a combination of the underlying system and the macrostate definitions; it is not a result of the MSM or haMSM analysis. We also examined how the presence of the plateau regions is affected by changing the core-likeness of the macrostates (Figure 17). The plateaus sharply disappear for all systems if highly core-like macrostates are used, while they are largely unaffected for small sizes of the intermediate regions.

Analogous data based on the kinetically-clustered macrostates (Figure 18) yields similar results for two of the systems (Trp-cage and NTL9), while for chignolin and villin the MFPTs are underestimated compared to PCCA++ results at short lag times and are missing the plateau regions. This suggests macrostates may not be ideally defined in the latter cases via kinetic clustering.

### Observable: Pathways/mechanism

Understanding mechanism is a key goal of molecular dynamics studies. In principle, both MSMs and haMSMs may be used to model transition mechanisms, but because MSMs are validated for a particular Markovian lag time, caution must be exercised in analyzing transitions which may occur on shorter timescales. As shown in Table 2, *t_T P_* is typically ≲ 10 ns for the protein transitions of interest, considerably less than the validated MSM lag times ≳ 100ns. For the sake of comparison and because lag time sensitivity will be of considerable interest, MSMs are here considered at a range of lag times, including values well below the validated lag times.

We analyze mechanism using three approaches that vary in “resolution” but all provide objective yardsticks for comparing models with one another and against empirical MD data.
i. The crudest measure simply tracks the fraction of direct transitions, defined to be those where the *microstate-discretized* macrostate-to-macrostate transition occurs without visiting *any* intermediate microstate; this statistic will depend on the model lag time and also the size of the intermediate region.
ii. Second, we employ an analysis based on the *configurational* distribution of the transition path ensemble, as described by Hummer,^60^ which aggregates transition paths together; although temporal information is removed, the resulting population profile over intermediate microstates provides a configurational representation of mechanism. A simple measure of some of the temporal information missing from the configurational representation is the average transition path length shown in the SI (Figure 19).
iii. Finally, employing time-sequential conformational information, we use the recently proposed “pathway histogram analysis of trajectories” (PHAT) method^87^ which classifies MD or model trajectories into pathway classes, yielding a path histogram which is a mechanistic signature of the transition. Classification in the PHAT approach is performed using the “fundamental sequence” (FS) of each transition trajectory; the FS, roughly, is the “backbone” of the transition path with loops and back-and-forth steps removed, expressed as a sequence of microstates traversed.^87^ For all three analyses, we combine events in both directions to obtain better statistics exploiting the symmetry of forward and reverse mechanisms under equilibrium conditions. ^55^

Configurational analysis of mechanism is embodied in Figs. 6 and 7. The haMSM accurately reproduces the fraction of direct pathways seen in MD for all the proteins and all sizes of the intermediate region. MSMs do not reproduce the MD well in general, except for some instances at very short lag times, which are well below the validated lag times *τ* ≳ 100 ns. Likewise, the haMSMs exhibit relatively low error by comparison to MD for *p*(*x*|*TP*_ind_), which is the configurational distribution over discrete microstates for the transition-path ensemble; the “ind” subscript indicates direct pathways probed in Figure 7 have been removed from the analysis. For *p*(*x*|*TP*_ind_), the MSMs perform best at short lag times well below the validated values.

**Figure 6:**
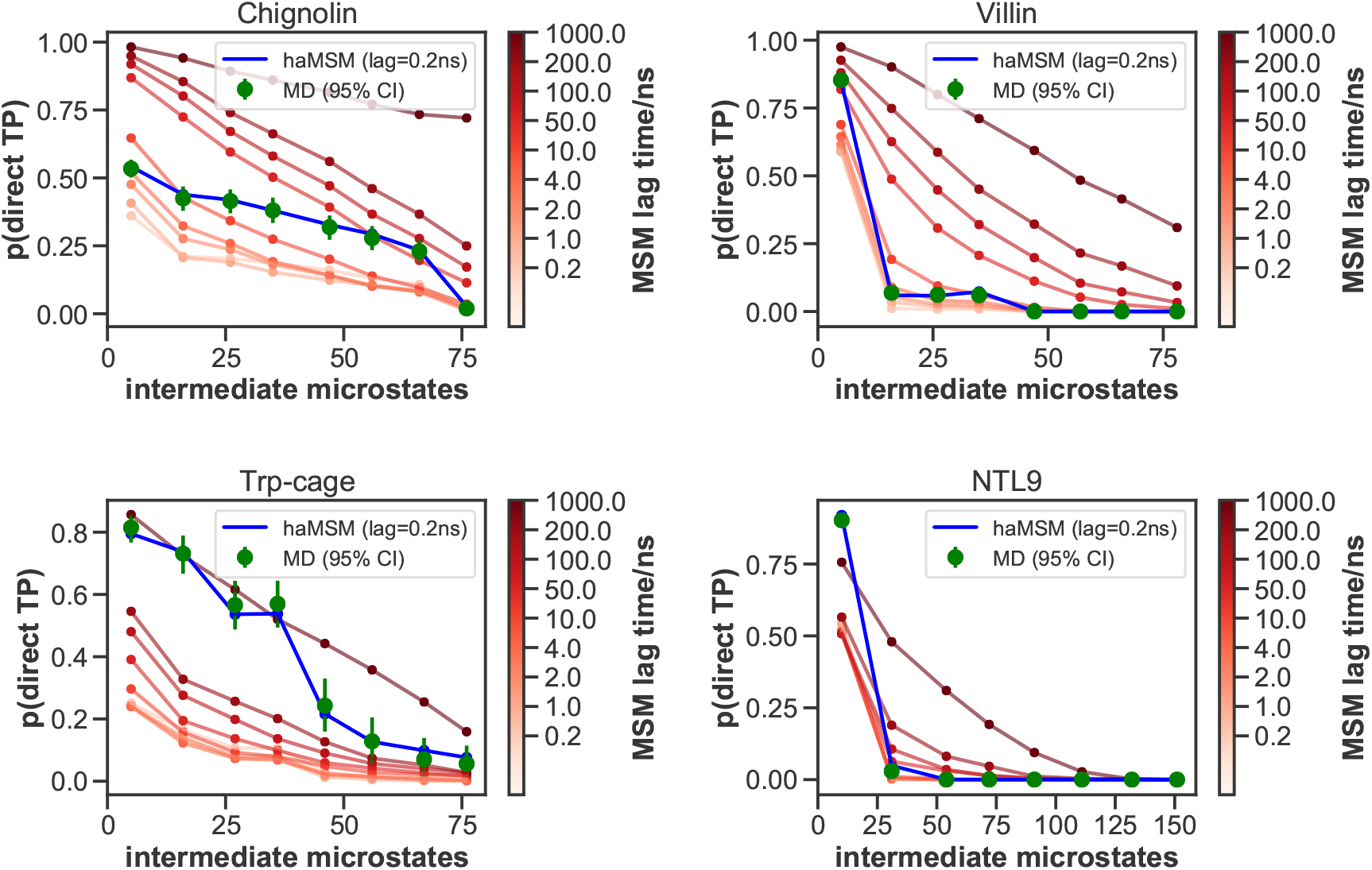
Simple mechanism comparison of MD, MSMs, and haMSMs using the fraction of direct folding pathways. For the given number of intermediate microstates, ensembles of discretized transition trajectories were analyzed to determine the fraction which directly ‘hopped over’ the intermediate region based on either the MD discretization time Δ*t*, also used for haMSM modeling, or else the indicated MSM lag time. A greater number of intermediate microstates indicates relatively smaller macrostates and accounts for the monotonic decrease of direct transitions.

**Figure 7:**
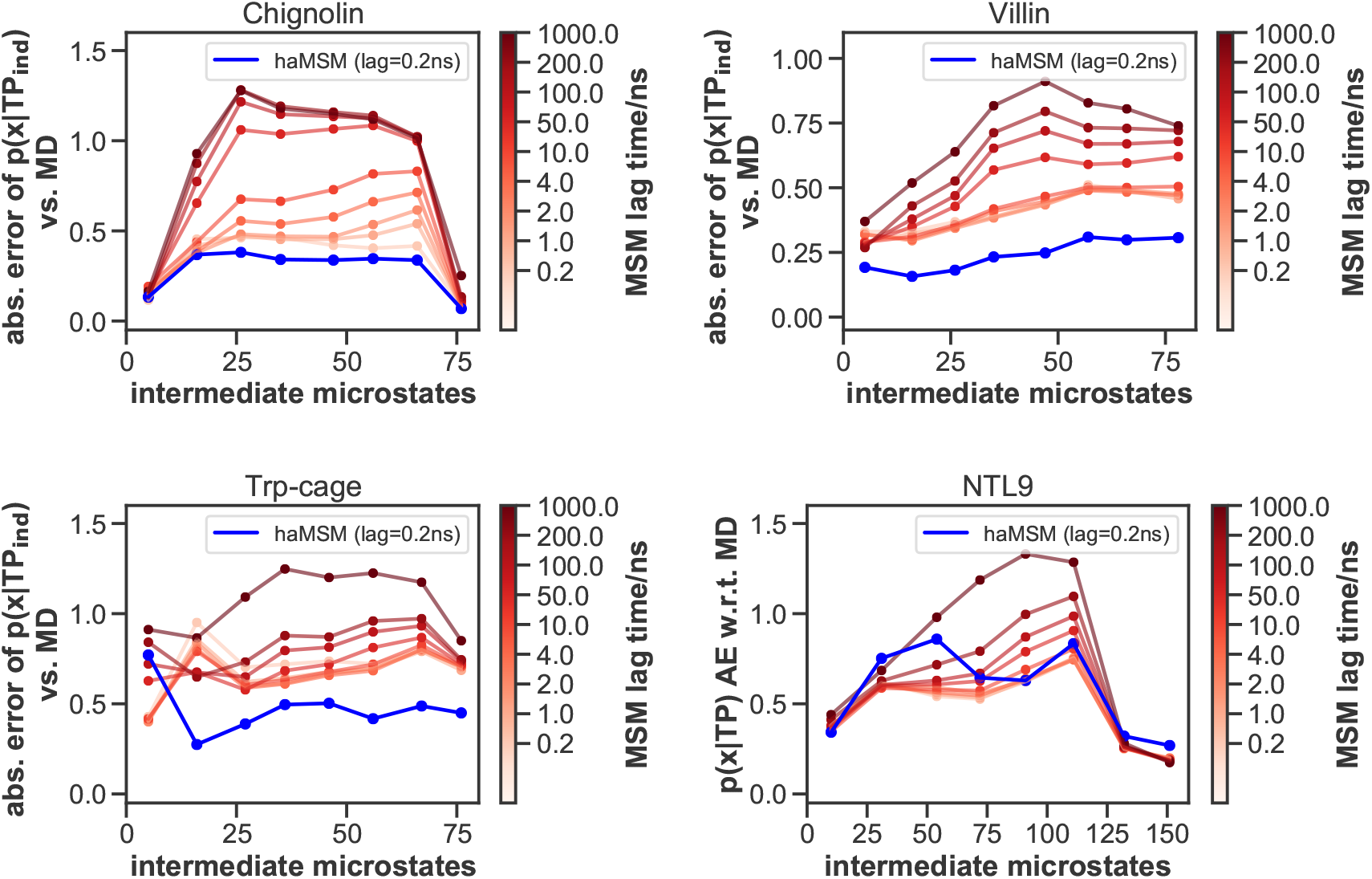
Mechanism comparison of MSMs and haMSMs to MD using the configurational distributions of transition path ensemble. Each panel plots the summed *absolute error, as compared to MD,* for intermediate microstate probabilities calculated for the transition path ensembles, i.e., *p*(*x|TP*_ind_), for the given number of intermediate microstates. Importantly, the “ind” subscript indicates that direct pathways analyzed in Figure 6 were excluded from the ensembles prior to computation of errors; had they been included, the MSM errors would be substantially larger.

Figure 8 shows the configurational analysis for a set number of intermediate microstates (10% of all states) and Figure 9 shows the comparison of MSMs and haMSMs with reference to pathway histogram data generated from MD using the same size of the intermediate. In both cases, the haMSMs recapitulate the configurational and mechanistic distributions found in long MD trajectories, and are largely successful even when only a small amount of history (1 or 10 ns) is used. The MSMs, however, exhibit irregular agreement with MD reference results: for most of the systems, no single lag time provides uniform agreement, while predictions for some of the microstates and for the most probable paths substantially differ from MD for many lags. The data also suggest that a fairly small number of pathway classes dominate the ensembles even though the number of intermediate microstates used for the analysis implies a large number of (mathematically) possible pathways.

**Figure 8:**
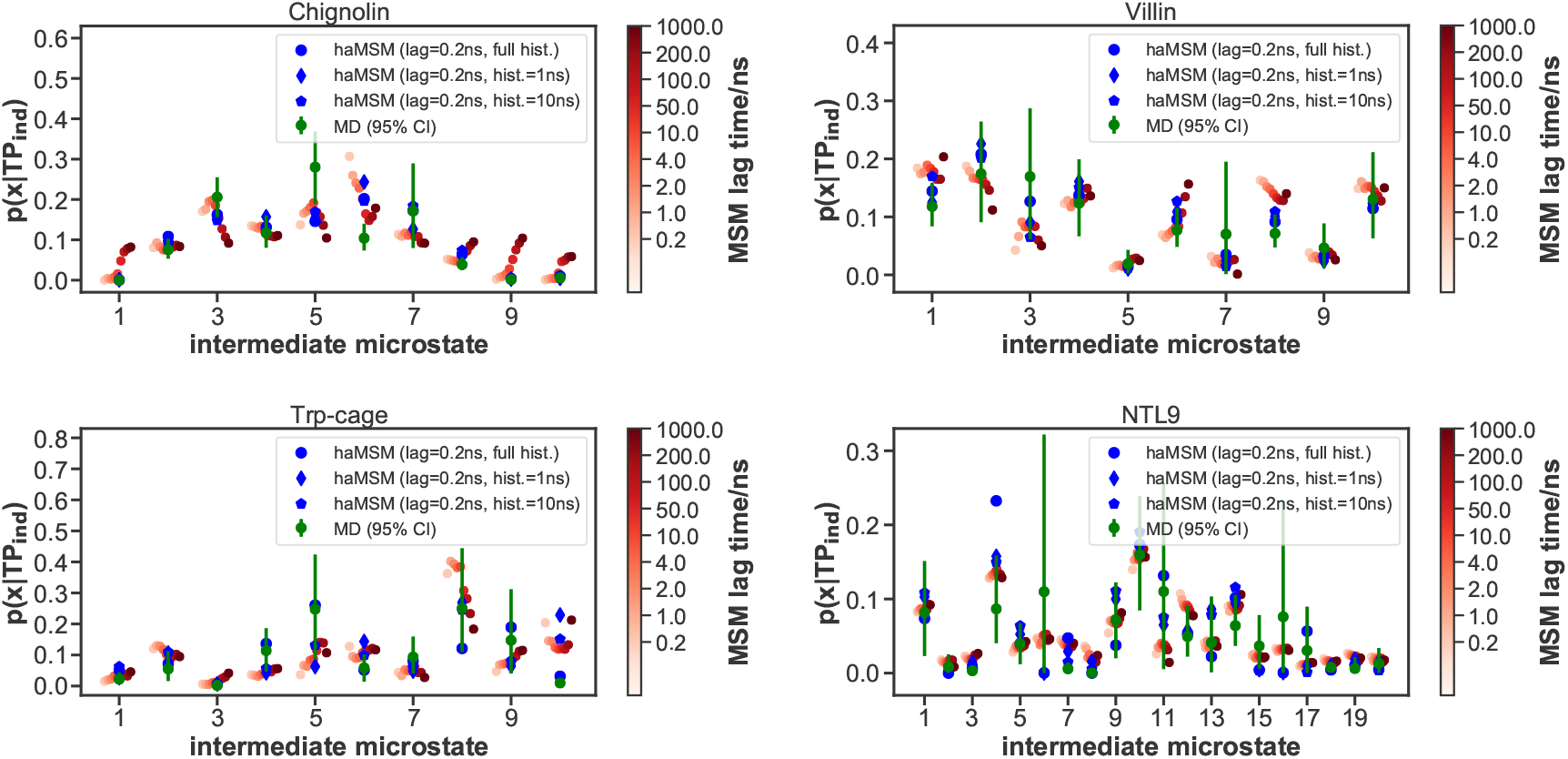
Mechanism comparison of MSMs and haMSMs to MD using the configurational distributions of transition path ensemble, for 10% intermediate states. For each protein, the probability distribution is plotted for different states to be on a transition pathway, i.e., *p*(*x|TP*_ind_), Importantly, the “ind” subscript indicates that direct pathways analyzed in Figure 6 were excluded from the ensembles prior to computation of errors; had they been included, the MSM errors would be substantially larger. The reference MD values (green) may be compared with MSM predictions for different lag times (red color scale at right) and haMSM estimates based on different amounts of history (blue symbols).

**Figure 9:**
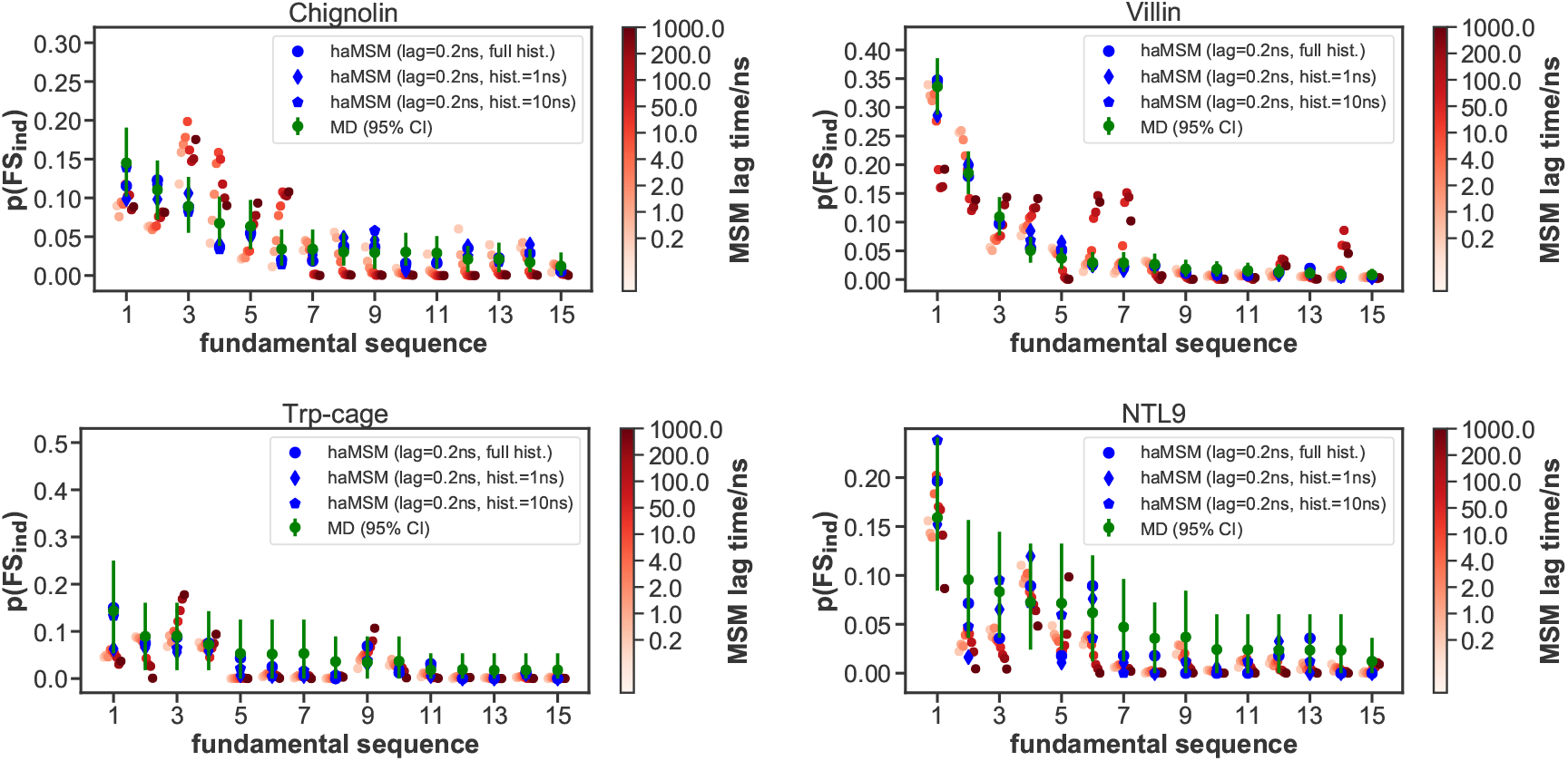
Path-based mechanistic comparison among MSMs, haMSMs, and MD. For each protein, the probability distribution is plotted for different mechanistic pathways based on the fundamental sequence (FS) approach. ^87^ The reference MD values (green) may be compared with MSM predictions for different lag times (red color scale at right) and haMSM estimates based on different amounts of history (blue symbols). Pathway indices are ordered based on decreasing probability in the MD reference data set. Only the top 15 paths are included. For the data shown, 10% of states were used as the intermediate to yield a manageable number of transition paths.

## Discussion and recommendations

Our study has filled an important gap in the MSM literature by direct and quantitative comparison of MSMs to the underlying long-time MD trajectories, both in terms of rate-constants for specific processes and mechanisms of protein folding. Prior studies typically examined implied timescales (ITS), which can be difficult to assign to specific structural transitions of interest, and did not characterize mechanisms in a way that enabled direct, quantitative comparison of MSMs with MD data. Furthermore, the MSMs employed in this study are among the most carefully validated in the literature, not only because of the size of the trajectory data sets^39^ used, but also due to the application of recently developed strict validation criteria. ^33,34^

Most centrally, we find for the folding and unfolding processes examined here, that:

i. validated MSMs provide reliable kinetics estimates at suitable lag times, contrary to what was implied by a recent analysis examining only the shortest lag times, ^72^ (ii) the lag times necessary for validated MSMs are too long to permit the detailed examination of transition events or to make mechanistic inferences, consistent with previous theoretical arguments,^18,36,49^ and (iii) augmenting MSMs using history information^72^ enables accurate kinetic analysis at short lag times and also yields mechanistic descriptions in quantitative agreement with MD.

The capabilities and limitations of MSMs stem directly from their mathematical basis. The validated MSM is constructed to match the eigenspectrum and time-correlation functions but not path-like properties relying on states with lifetimes shorter than the lag-time. The first-hit characteristic of the MFPT, which in a sense is path-dependent, likely disrupts agreement with MSMs at short lag times, whereas at longer lags the MFPT presumably behaves more like a correlation time in harmony with the MSM’s eigenspectrum. The haMSM’s construction explicitly accounts for the macrostate-to-macrostate directionality and appears to provide a reasonable approximation to path-like quantities, including MFPTs at short lag times.

Our findings raise several issues. First, in practical terms, what lag times should users expect will be needed in other systems and how does that affect the strategy for collecting MD data for MSM construction? We examined a series of relatively small, single-domain proteins based on effectively exhaustive MD sampling. ^39^ In more complex systems, our data suggest lag times exceeding 100 ns should be expected, and accordingly continuous trajectories on the *μ*s scale would be advisable. We advise users to examine ITS behavior as a function of lag time on both log and linear scales, because the logarithmic scale can be deceptive in suggesting a plateau when ITS values may still be increasing.

Our finding that the validated lag times for MSMs exceed typical transition path times (event durations) is cautionary. Users primarily interested in deriving mechanistic insights may want to pursue tools beyond standard MSMs. Mechanistic conclusions in older MSMs based on less complete validation may warrant re-examination.

For history-augmented MSMs, the present study suggests that including ≲ 50 ns of trajectory history is sufficient for estimating kinetic and mechanistic observables, pointing to the value of continuous trajectories exceeding 100 ns, consistent with prior work on first-passage times.^72^ Once a sufficient amount of history is included in the haMSM analysis, arbitrarily small time-discretizations (lag times) can be examined reliably for both kinetics and mechanism.

What accounts for the success of haMSMs in predicting mechanism quantitatively, despite that they are exact only for the *mean* FPT and approximate for other non-equilibrium observables? The haMSM transition matrix is built from transition counts in the *subset* of A-to-B directed (*α*) trajectories, and so is constructed to mimic the “forward” tendency embodied in that ensemble. To the extent that the distribution of mechanisms is predicted quantitatively, this means that the average transition probabilities in the *α* ensemble are not significantly different from the detailed tendencies which would be embodied, for instance, in a higher-degree Markov model conditioned on an extended sequence of prior states. Some physical intuition can come from a toy example where there are two transition pathways separated from each other by an energy barrier—i.e., a ridge. So long as none of the microstates (which are coarse-grained regions in configuration space) straddle the ridge, we would not expect a significant difference between the haMSM paths and the true paths. Any microstate straddling the ridge could, however, lead to unphysical crossover between the pathways; evidently, this latter occurrence is infrequent in the systems and models examined here. Finally, note that haMSMs exhibited similar success in reproducing the *distribution* of FPTs, which also is only predicted approximately.^43^

On the whole, we hope these findings provide guidance for users of MSMs and haMSMs, though we acknowledge that additional similar comparisons for different types of processes, such as conformational changes and ligand binding, as well as more complex systems, would be of great value for the community.

## Footnotes

1 Google Scholar search: ‘“markov-state-models” molecular dynamics’

2 The haMSMs considered here include history information that is not part of a system’s standard phase-space description and thus were termed ‘non-Markovian’ in prior reports. ^41–43^ However, it should be noted that these models can formally be written as Markov models with history information encoded as an auxiliary variable.

3 More advanced approaches to MSM construction involve the use of *core sets*, ^26,29,50,51^ described in more detail below.

## Acknowledgments

We thank D. E. Shaw Research (DESRES) for providing a copy of the protein folding trajectory data set from. ^39^ Simon Olsson helped to build early MSM models for this study. DMZ acknowledges support from NIH Grant R01GM115805, as well as NSF Grant MCB-1119091. JDC acknowledges support from NIH Grant R01GM121505 and National Cancer Institute Cancer Center Core grant P30CA008748. RPW acknowledges support from the Tri-Institutional PhD Program in Chemical Biology and the Department of Defense (Peer Reviewed Cancer Research Program, Award W81XWH-17-1-0412). This project has also been funded in part with federal funds from the National Cancer Institute, National Institutes of Health, under contract HHSN26120080001E. The content of this publication does not necessarily reflect the views or policies of the Department of Health and Human Services, nor does the mention of trade names, commercial products, or organizations imply endorsement by the U.S. Government. We thank Josh Fass for helpful discussions.

## Author Contributions

Conceptualization, E.S., J.D.C. and D.M.Z.; Methodology, E.S., R.P.W, S.O., C.W., J.D.C., and D.M.Z; Software, E.S., R.P.W., S.O., and C.W.; Investigation, E.S., R.P.W., and D.M.Z.; Writing - Original Draft, E.S., R.P.W., J.D.C., and D.M.Z.; Writing - Review & Editing, E.S., R.P.W., F.N., J.D.C., and D.M.Z.; Visualization, E.S. and R.P.W.; Supervision, F.N., J.D.C., and D.M.Z.; Funding Acquisition, R.P.W., F.N., J.D.C., and D.M.Z.

## Disclosures

JDC was a member of the Scientific Advisory Board for Schrödinger, LLC during part of this study, and is a current member of the Scientific Advisory Board of OpenEye Scientific Software. The Chodera laboratory receives or has received funding from multiple sources, including the National Institutes of Health, the National Science Foundation, the Parker Institute for Cancer Immunotherapy, Relay Therapeutics, Entasis Therapeutics, Silicon Therapeutics, EMD Serono (Merck KGaA), AstraZeneca, Vir Biotechnology, XtalPi, the Molecular Sciences Software Institute, the Starr Cancer Consortium, the Open Force Field Consortium, Cycle for Survival, a Louis V. Gerstner Young Investigator Award, and the Sloan Kettering Institute. A complete funding history for the Chodera lab can be found at http://choderalab.org/funding.

## Supporting Information

### MSM scoring

MSMs at a lag time of 100 ns were constructed using discrete microstate trajectories from the training set and scored on the test set trajectories. For comparison, we also performed scoring at a short MSM lag time of 10 ns, hypothesizing this could better optimize the reproduction of kinetics at short lag times for comparison with haMSMs. We used the 100 ns lag time top models for all analysis, and compared the results to the 10 ns lag time models in SI figures.

We used a 50:50 shuffle-split cross-validation scheme to find the optimal set of hyperparameters while avoiding overfitting. In this scheme, 2 *μ*s long fragments of the trajectories (i.e., the original fragments in which the datasets are provided by DESRES) are randomly split into training and test sets of approximately equal sizes. To obtain standard deviations indicative of out-of-sample model performance, this shuffle-split model evaluation procedure was repeated 10 times with different random divisions of the dataset into training and test sets. Scoring was based on the sum of squared-eigenvalues of the transition matrix (VAMP-2 score^32^), as this particular score is physically interpretable as ‘kinetic content’.

To choose the best number of top eigenvalues to score the models with, we initially performed scoring separately at each number of top eigenvalues between 2 and 50. We then chose the number of eigenvalues for which the score of the top scoring model was closest to 50% of the number of eigenvalues (i.e., the highest possible score), in order to maximize the signal from the true dynamical processes and increase the resolution of the scores, while minimizing the noise from spurious eigenvalues (Figure 10, Figure 11 shows the analogical results for the 10 ns scoring lag time). Hence 3 (4 at 10 ns) top eigenvalues were used for chignolin, 5 (12 at 10 ns) eigenvalues for villin, 6 (17 at 10 ns) eigenvalues for Trp-cage, and 5 (38 at 10 ns) eigenvalues for NTL9. To evaluate a large set of hyperparameters, reduced datasets subsampled to 10 ns/frame (for NTL9: 10 ns/frame at 10 ns scoring lag time, increased to 50 ns/frame at 100 ns scoring lag time) were used for computational feasibility, except for chignolin, which remained at 0.2 ns/frame intervals due to its small size. The datasets were featurized with all minimal residue–residue distances (calculated as the closest distance between the heavy atoms of two residues separated in sequence by at least two neighboring residues). For consistency in interpretation and computational feasibility, this featurization choice was made without variational scoring. The datasets were projected into a kinetically relevant space using tICA,^40,64^ at lag time 10 ns (50 ns for NTL9 at the 100 ns MSM scoring lag due to higher subsampling; for chignolin lag times 1 ns and 5 ns were also possible due to no subsampling), with either kinetic ^65^ or commute^66^ mapping, retaining the following numbers of tICs (maximum number of which depends on the number of features and hence size of the protein): 2 or the number of tICs corresponding to 95% of total kinetic variance/content (”95%”) for chignolin; 2, 10, 50, 100, 150, or 95% for Trp-cage; 2, 10, 50, 100, 300, 500, or 95% for villin and NTL9. Each of the tICA outputs was discretized using k-means clustering into 50, 100, 300, 500, 800, or 1000 microstate clusters (except for chignolin, where 200, 400, 600, or 900 microstate clusters were also tried for better scoring resolution, due to the smaller number of features and hence number of tICs retained). Table 3 summarizes all hyperparameter options assessed.

**Figure 10:**
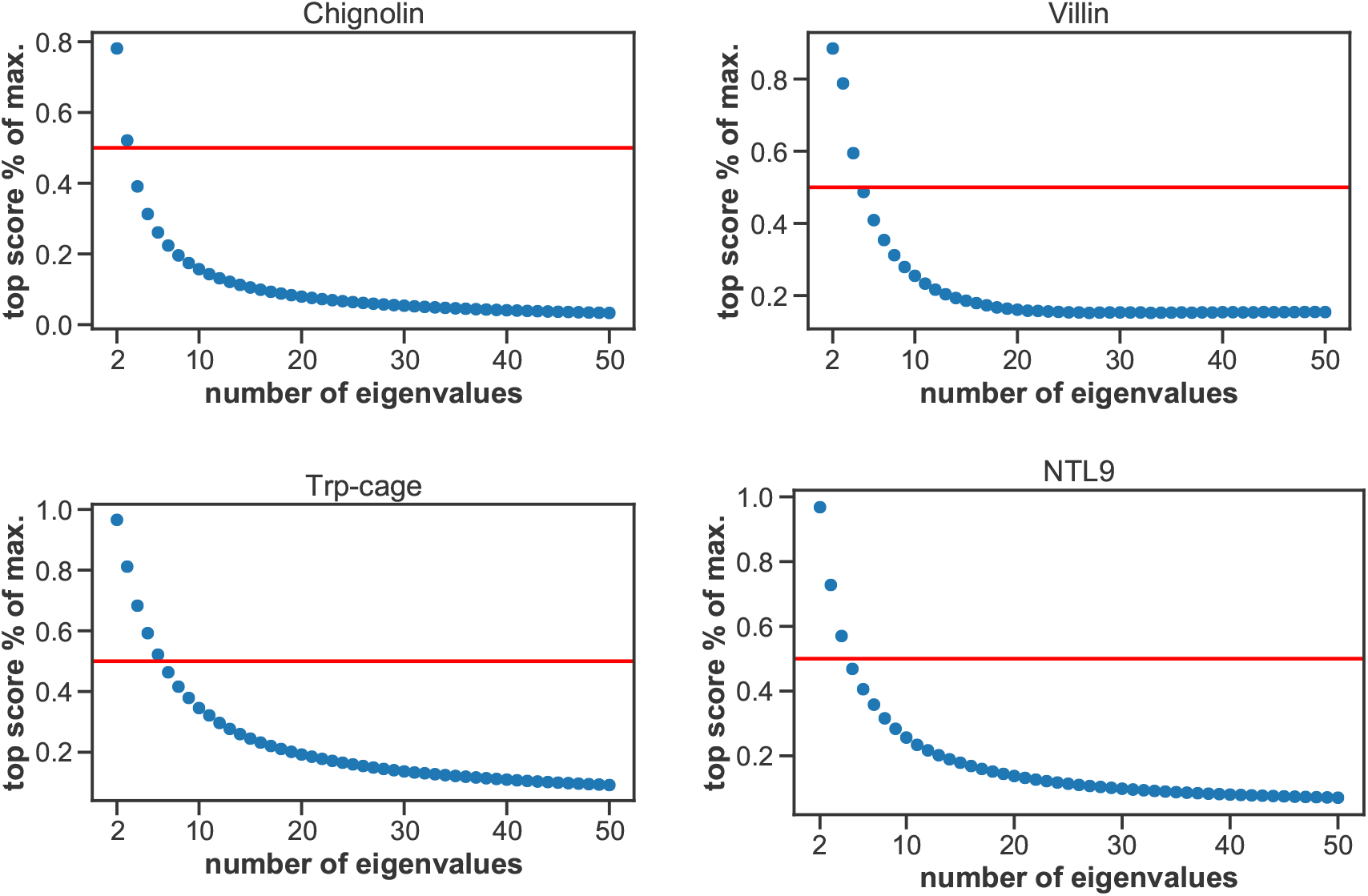
Selection of the number of top eigenvalues for scoring, at a 100 ns lag time. Markov state models (MSMs) were VAMP-2 scored separately at each number of top eigenvalues between 2 and 50 included in the scoring. The ratios of the top scoring models at each choice of numbers of eigenvalues and that number of eigenvalues (i.e. the highest possible score) are plotted. For selection of the final model, we chose the number of eigenvalues for which the ratio was closest to 0.5, marked by the red horizontal line.

**Figure 11:**
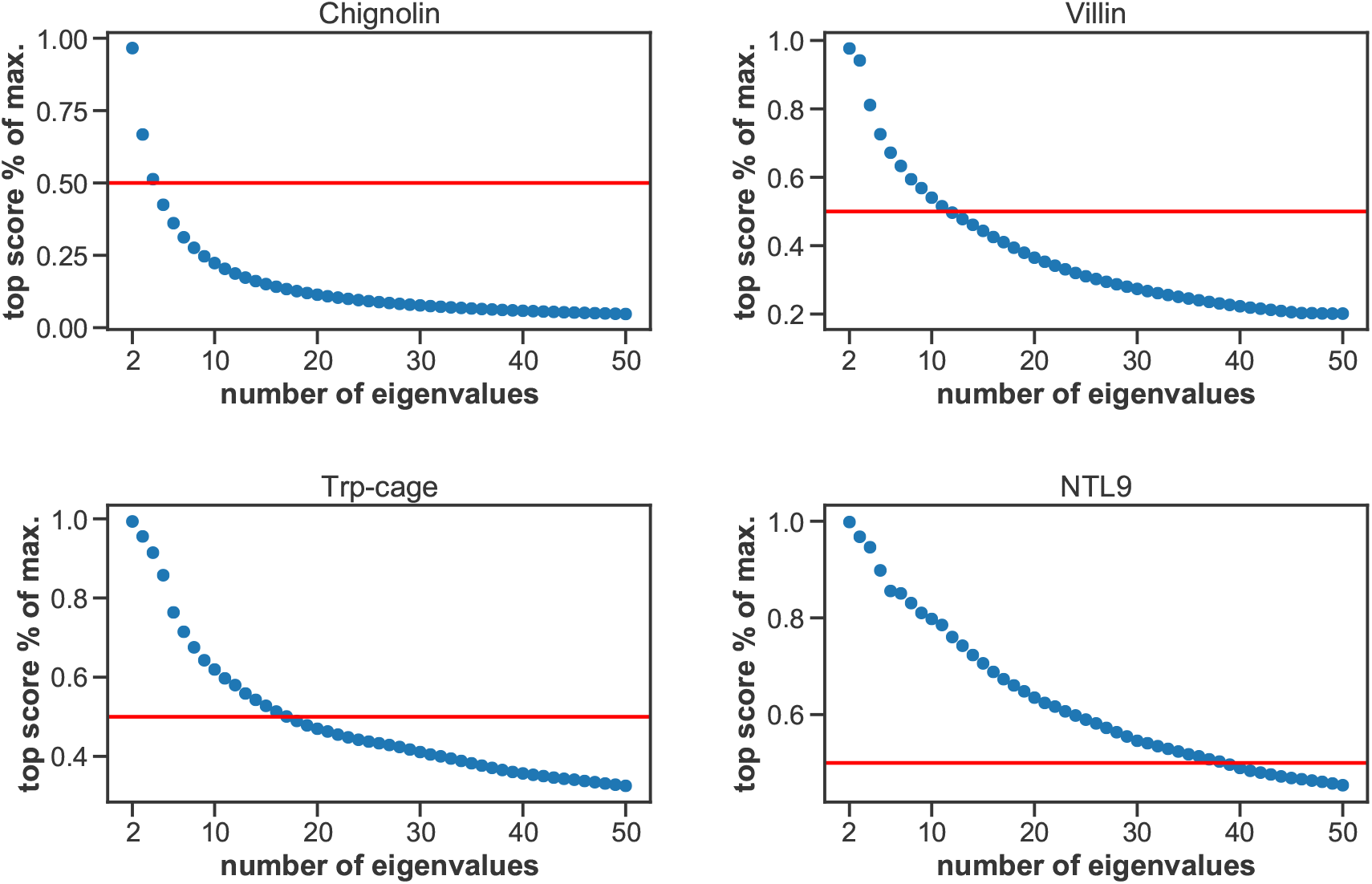
Selection of the number of top eigenvalues for scoring, at a 10 ns lag time. Markov state models (MSMs) were VAMP-2 scored separately at each number of top eigenvalues between 2 and 50 included in the scoring. The ratios of the top scoring models at each choice of numbers of eigenvalues and that number of eigenvalues (i.e. the highest possible score) are plotted. For selection of the final model, we chose the number of eigenvalues for which the ratio was closest to 0.5, marked by the red horizontal line.

**Figure 12:**
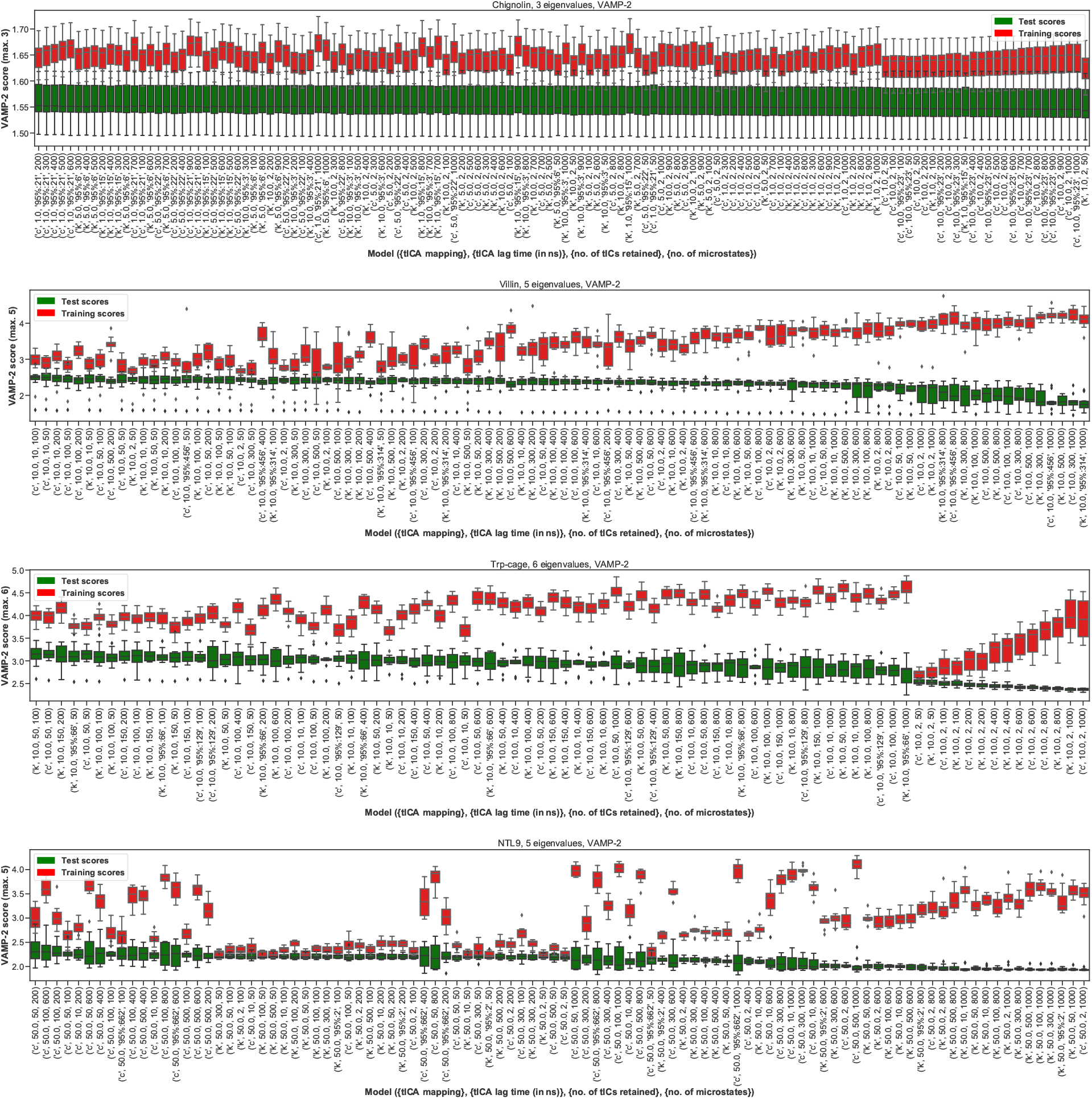
VAMP-2 scoring results for optimal hyperparameter choice, at a 100 ns lag time. The distributions of the VAMP-2 scores of ten shuffle-splits of the data for each individual set of hyperparameters (model) are shown as box-and-whisker plots. Bands of boxes show the first, second, and third quartiles, while whisker ends represent the lowest and highest scores still within 1.5 of the interquartile range from the first and third quartiles respectively. Scores lying outside of that range are shown as diamonds. The models are denoted as ([tICA mapping], [tICA lag time (in ns)], [number of tICs retained], [number of microstates]). Test scores are shown in green and training scores in red.

Figure 12 shows the results of the scoring at the 100 ns lag time, and Figure 13 at the 10 ns lag time. Model scores are reported below as means with standard deviations over 10 shuffle-splits. As there were no statistically significant differences between the scores for chignolin at the 100 ns lag time, we used the top model scored at 10 ns for all analyses. We also note the top scoring models for villin were identical at both lag times.

**Figure 13:**
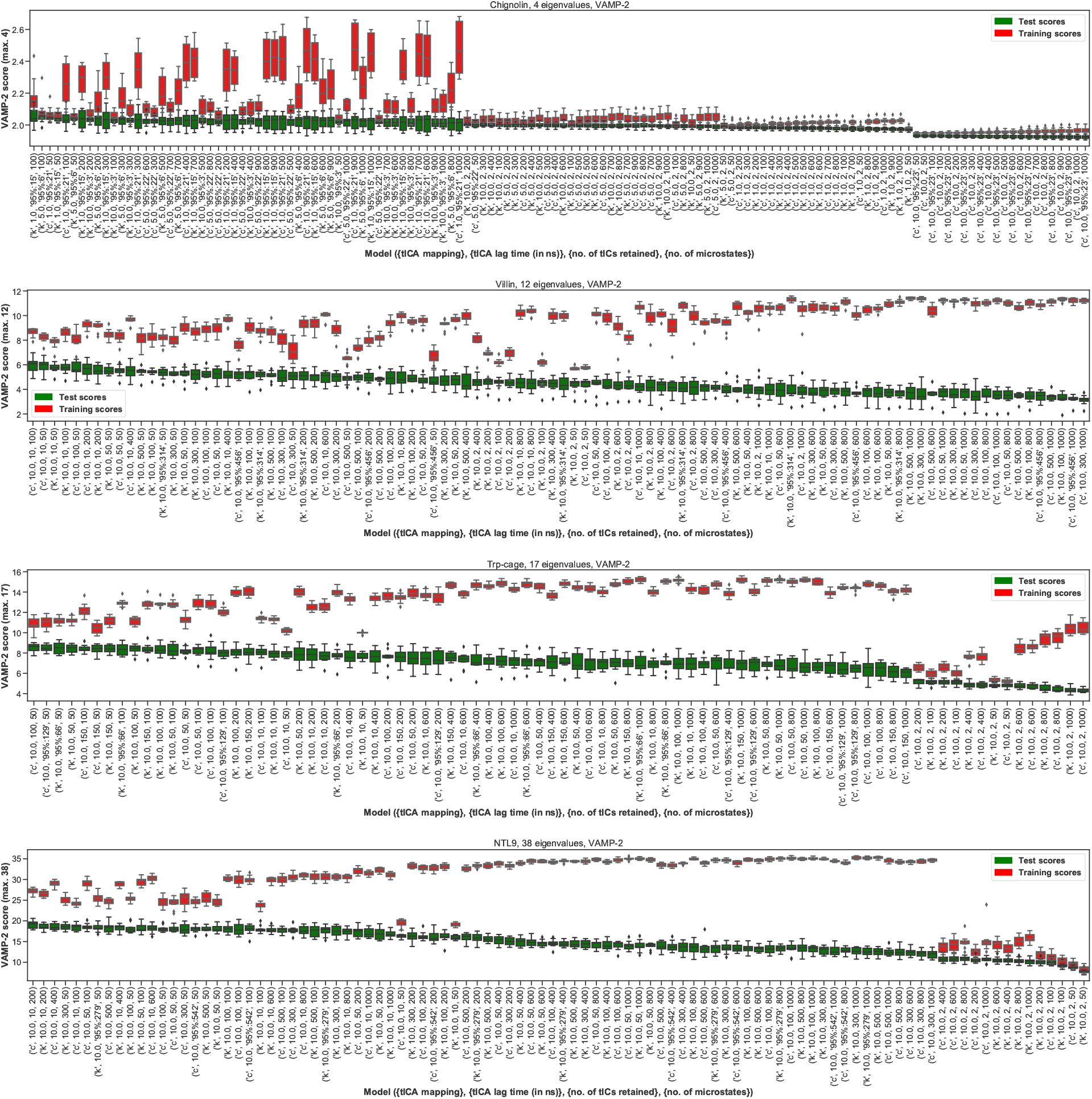
VAMP-2 scoring results for optimal hyperparameter choice, at a 10 ns lag time. The distributions of the VAMP-2 scores of ten shuffle-splits of the data for each individual set of hyperparameters (model) are shown as box-and-whisker plots. Bands of boxes show the first, second, and third quartiles, while whisker ends represent the lowest and highest scores still within 1.5 of the interquartile range from the first and third quartiles respectively. Scores lying outside of that range are shown as diamonds. The models are denoted as ([tICA mapping], [tICA lag time (in ns)], [number of tICs retained], [number of microstates]). Test scores are shown in green and training scores in red.

**Figure 14:**
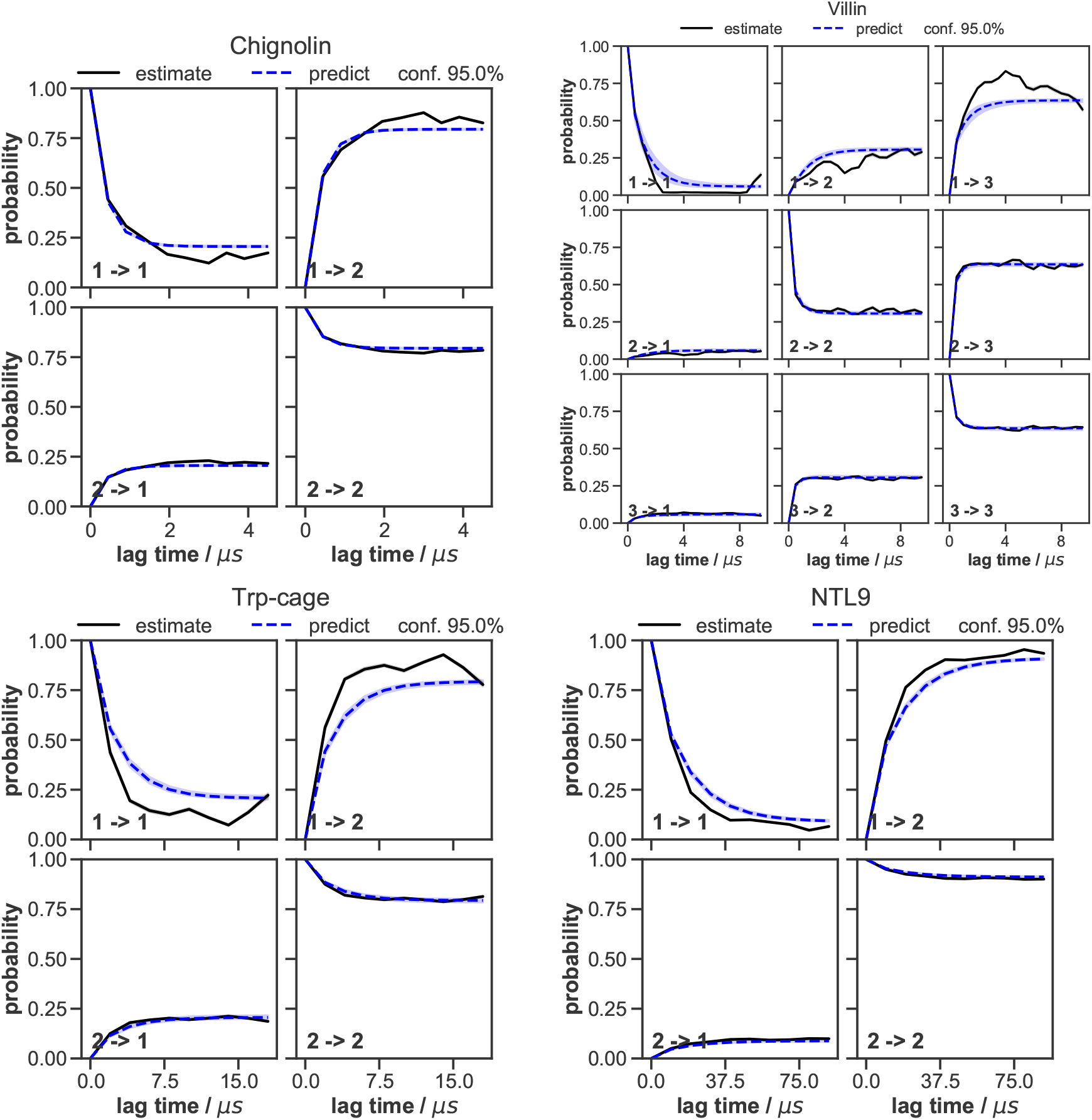
Chapman-Kolmogorov tests of the Bayesian Markov state models constructed for each system. The objective of the Chapman-Kolmogorov test is to assess the kinetic self-consistency of the MSM, i.e., whether the predictions of longer time behavior made from the BMSM being tested match the estimates made from BMSMs generated at longer lag times. For each macrostate, probability density is assigned to the BMSM microstates according to their metastable memberships to the given macrostate and evolution of the probability in time in the tested BMSM is plotted in blue (”predictions”). At those same longer lag times new BMSMs are estimated and their probability densities of being in the given macrostate after one lag time are plotted in black (”estimates”). The shaded regions correspond to the 95% confidence intervals of the mean of the predictions and estimates (the estimate confidence intervals are very narrow).

The following top scoring models were selected using the 100 ns scoring lag time: chignolin: all models statistically the same; villin: commute tICA mapping, 10 ns tICA lag time, 10 tICs, 100 microstates, score 2.43 (5 eigenvalues, SD: 0.31); Trp-cage: kinetic tICA mapping, 10 ns tICA lag time, 50 tICs, 100 microstates, score 3.13 (SD: 0.24, 6 eigenvalues); NTL9: commute tICA mapping, 50 ns tICA lag time, 50 tICs, 200 microstates, score 2.34 (SD: 0.26, 5 eigenvalues).

The following top scoring models were selected using the 10 ns scoring lag time: chignolin: kinetic tICA mapping, 1 ns tICA lag time, 15 tICs (95% kinetic variance), 100 microstates, score 2.05 (4 eigenvalues, SD: 0.05); villin: commute tICA mapping, 10 ns tICA lag time, 10 tICs, 100 microstates, score 5.96 (12 eigenvalues, SD: 0.58); Trp-cage: commute tICA mapping, 10 ns tICA lag time, 100 tICs, 50 microstates, score 8.51 (SD: 0.42, 17 eigenvalues); NTL9: commute tICA mapping, 10 ns tICA lag time, 10 tICs, 200 microstates, score 19.11 (SD: 0.89, 38 eigenvalues).

Finally, to construct the discrete microstate trajectories used in this work, the modeling process was repeated with full datasets (with no additional striding, i.e. at 0.2 ns/frame intervals and with no train-test splitting) using the top scoring parameters for the repeated tICA and k-means calculations.

### Coarse-graining into macrostates PCCA++ villin coarse-graining

Our villin MSM identified a microsecond timescale (∼1.1 *μ*s), which was not present in a previously published MSM of this dataset,^81^ likely due to our use of the residue-residue distances featurization combined with tICA, compared to the minRMSD metric in.^81^ By coarse-graining the MSM into two macrostates, we identified this longer timescale as corresponding to the transition between the “folded – unfolded” and “misfolded” macrostates. The “misfolded” macrostate shows formation of short-lived helicity between residues ASN60 and LEU63 (Figures 15, 16). The previously^81^ identified folding timescale of ∼ 400 ns is the second slowest timescale in our MSM, and a 3 macrostate coarse-graining was necessary to obtain separation between the “folded” and “unfolded” macrostates. All folding kinetics are considered only between the “folded” and “unfolded” macrostates, with no regard to the “misfolded” macrostate.

**Figure 15:**
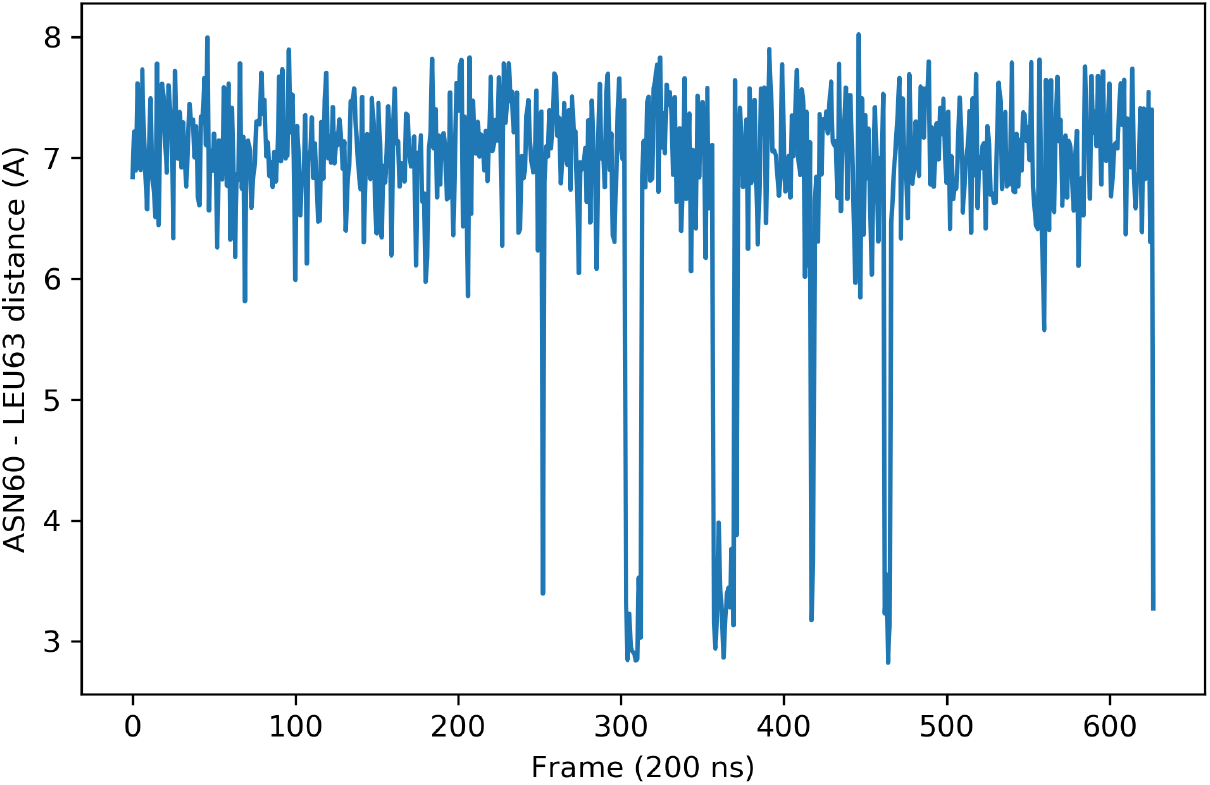
ASN60 - LEU63 minimum distance along the villin trajectory. Shortlived “misfolded” states can be seen, explaining the appearance of a very long timescale that does not correspond to folding in the MSM (see the ‘PCCA++ villin coarse-graining’ SI section for details).

**Figure 16:**
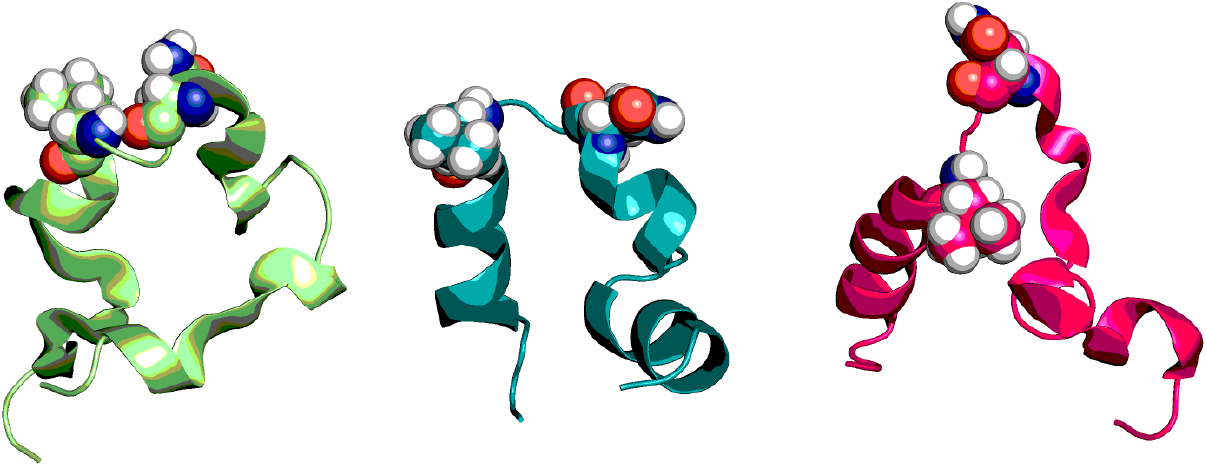
Sample frames from the three macrostates of the villin BMSM. From left to right, cartoon representations of “misfolded”, “folded”, and “unfolded” frames are shown. ASN60 and LEU63 are also shown in spheres. The presence of the “misfolded” states explains the appearance of a very long timescale that does not correspond to folding in the MSM (see the ‘PCCA++ villin coarse-graining’ SI section for details).

**Figure 17:**
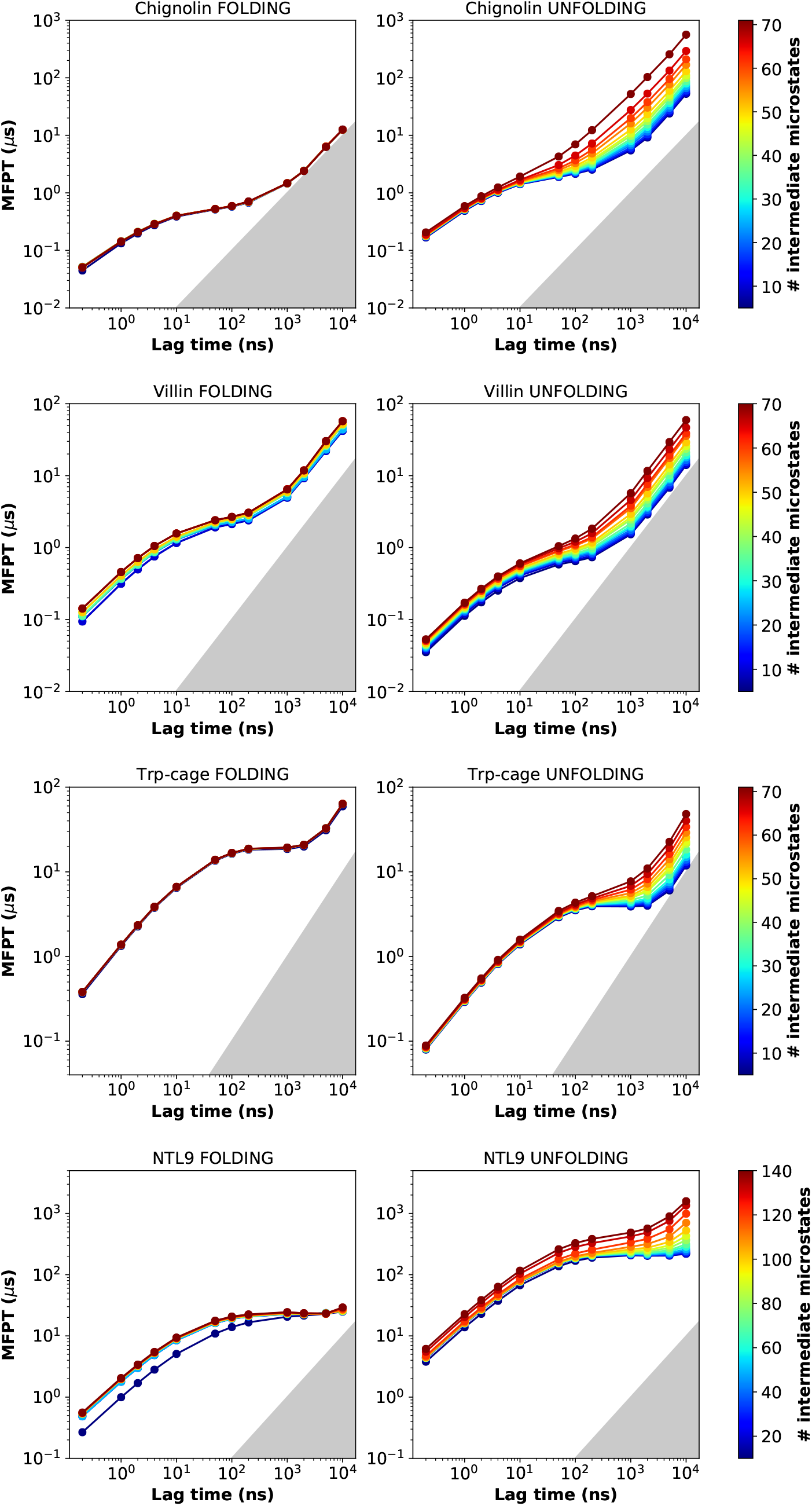
Sensitivity of MFPTs calculated from Markov state models to the core-likeness of the macrostates. The MFPT for both folding and unfolding calculated from MSMs is plotted as a function of lag time. The curves are colored by the number of microstates defined as the intermediate region - the larger the intermediate, the more core-like the macrostates become. The gray area signifies region where MFPTs become equal to or smaller than the lag time and can no longer be resolved.

**Figure 18:**
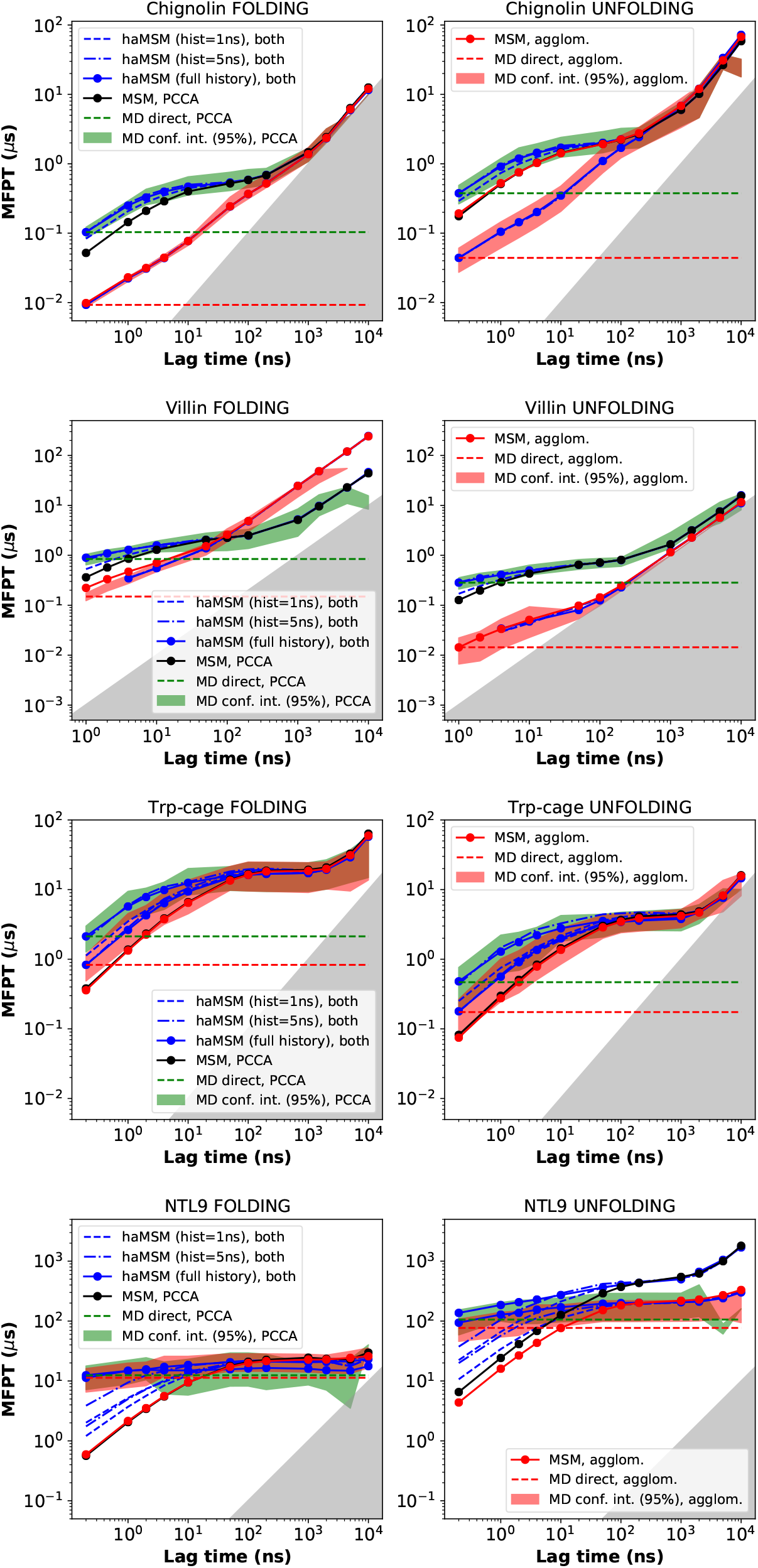
Comparison of the MFPT dependence on lag time for macrostates defined by PCCA++ or agglomerative clustering. The MFPT for both folding and unfolding is plotted as a function of lag time. Reference MD data is shown as the 95% confidence interval (green bands for PCCA a5n7d red bands for agglom.), which can be compared to validated MSM data (black lines for PCCA and red lines for agglom.) and haMSM values with full history (solid blue lines for both methods) and partial history (dashed blue lines for both methods). The PCCA macrostates are defined using a cutoff such that the

**Figure 19:**
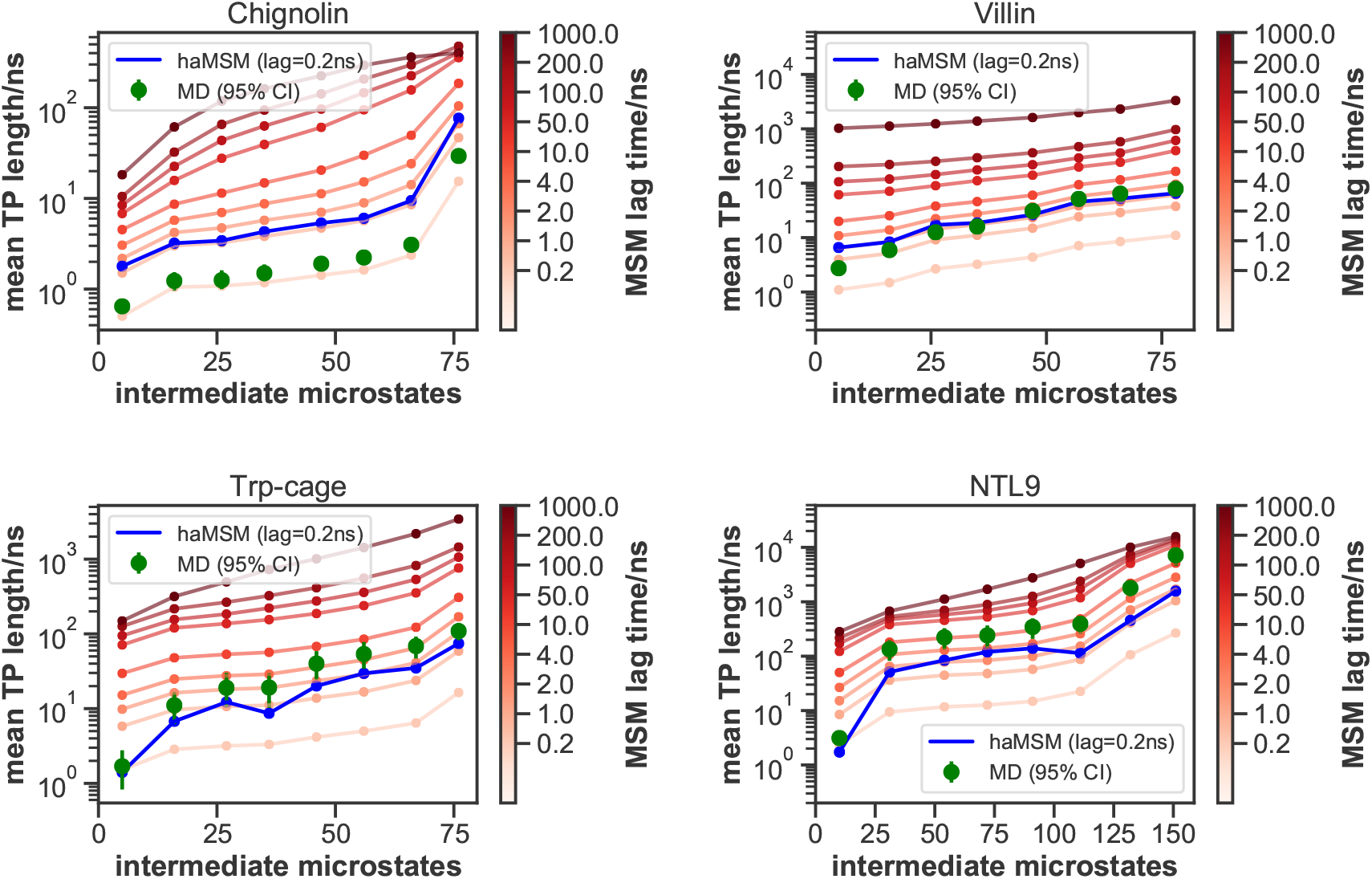
Simple mechanism comparison of MD, MSMs, and haMSMs using the mean lengths of transition paths. For the given number of intermediate microstates, ensembles of discretized transition trajectories were analyzed to determine the mean of the distribution of all transition paths. Differently from the configurational analysis of indirect paths, we include the direct paths here; the transition paths include the last frame in the origin macrostate and the first frame in the destination macrostate, i.e. the length of a direct path is 2 x the lag time.

**Figure 20:**
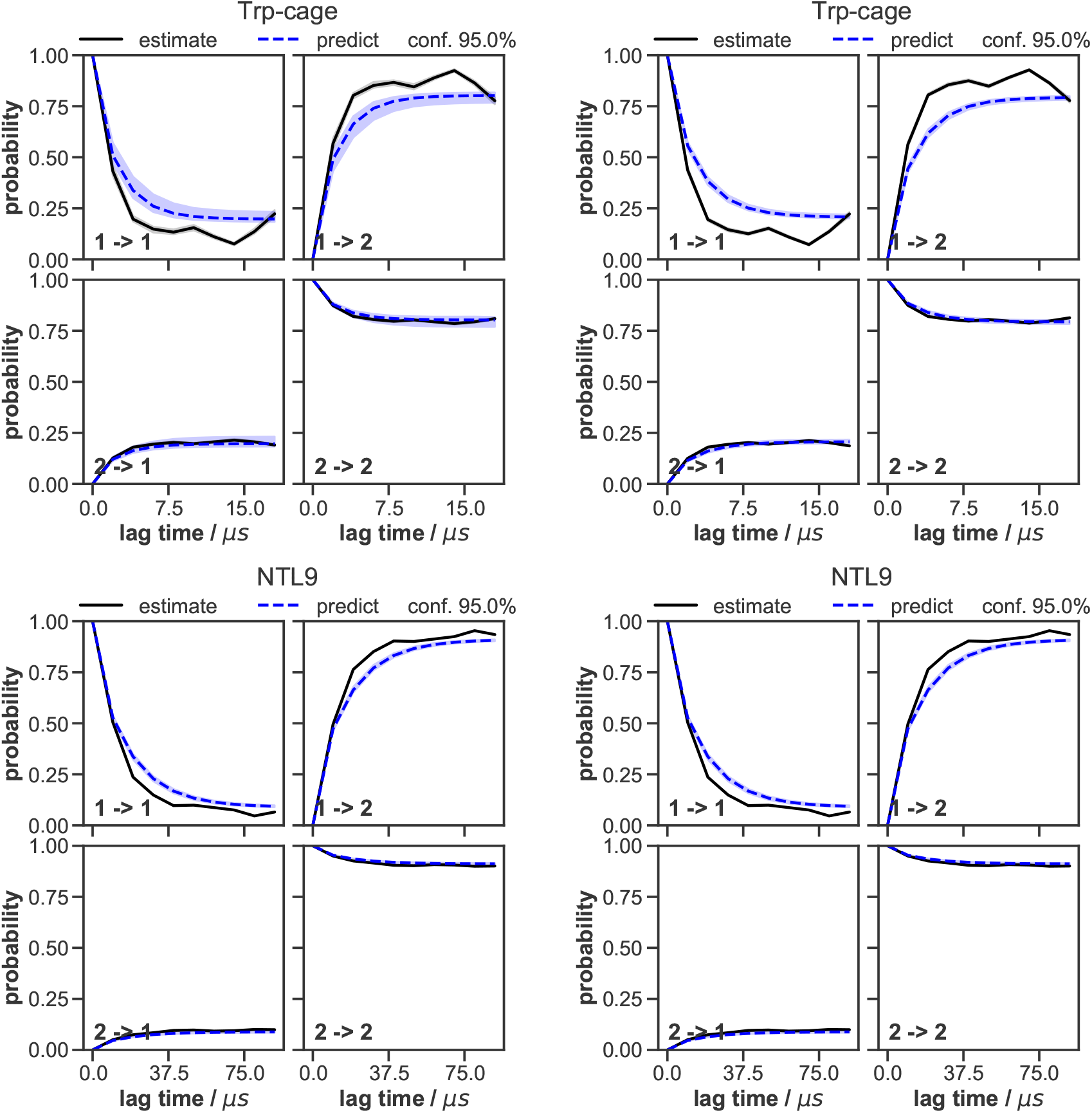
Comparison of top models selected at scoring lag times 100 ns (top) and 10 ns (bottom), for Trp-cage and NTL9: Chapman-Kolmogorov tests of the Bayesian Markov state models constructed for each system. The objective of the Chapman-Kolmogorov test is to assess the kinetic self-consistency of the MSM, i.e., whether the predictions of longer time behavior made from the BMSM being tested match the estimates made from BMSMs generated at longer lag times. For each macrostate, probability density is assigned to the BMSM microstates according to their metastable memberships to the given macrostate and evolution of the probability in time in the tested BMSM is plotted in blue (”predictions”). At those same longer lag times new BMSMs are estimated and their probability densities of being in the given macrostate after one lag time are plotted in black (”estimates”). The shaded regions correspond to the 95% confidence intervals of the mean of the predictions and estimates (the estimate confidence intervals are very narrow).

**Figure 21:**
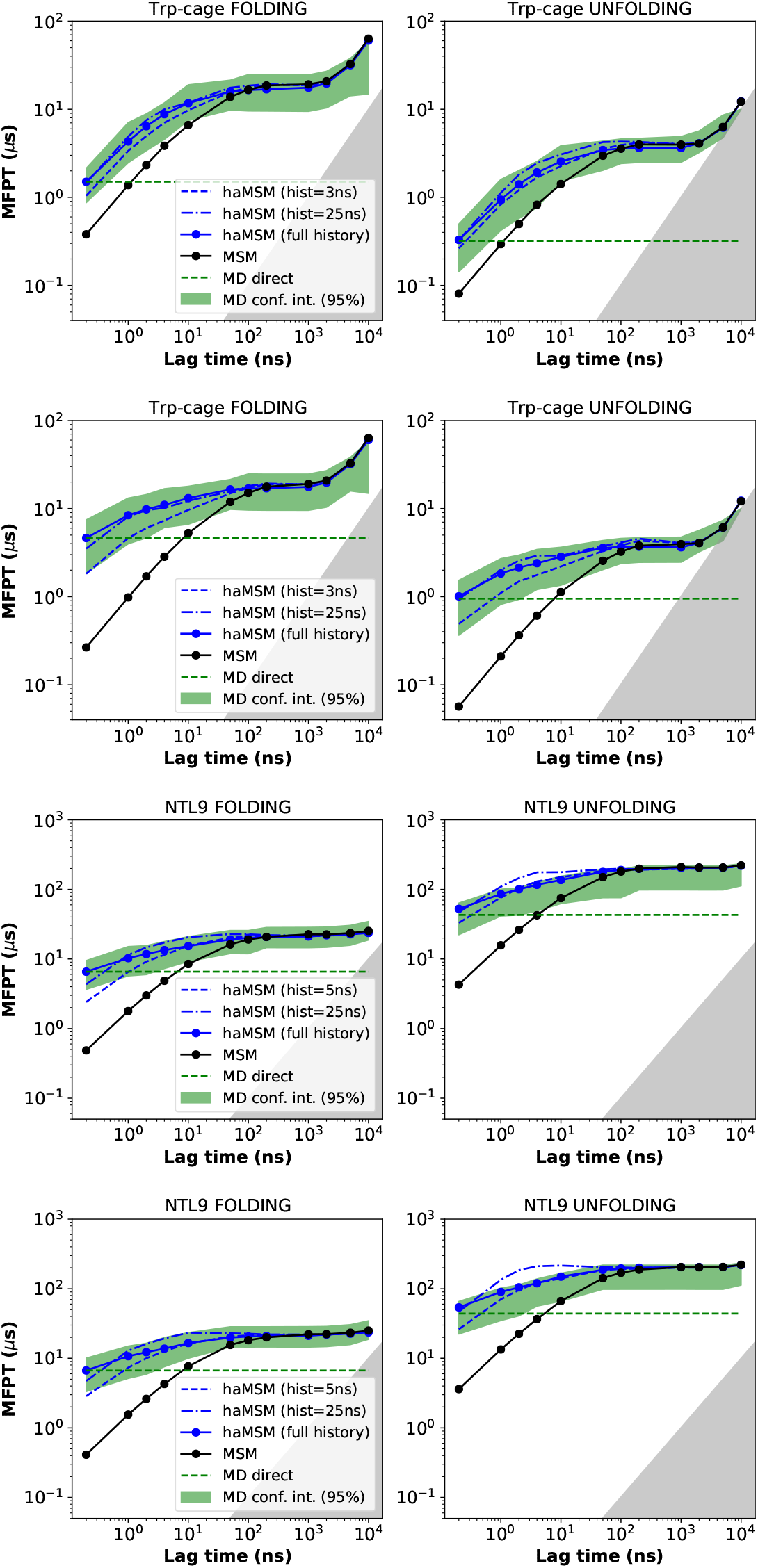
Comparison of top models selected at scoring lag times 100 ns (top) and 10 ns (bottom), for Trp-cage and NTL9: MFPT estimates compared among MD, MSMs, and haMSMs. The MFPT for both folding and unfolding is plotted as a function of lag time. Reference MD data is shown as the 95% confidence interval (green band), which can be compared to validated MSM data (black lines) and haMSM values with full history (solid blue lines) and partial history (dashed blue lines). The gray area signifies the region where MFPTs become equal to or smaller than the lag time and can no longer be resolved. The MD confidence intervals missing for the final data points of chignolin and villin are due to no more transition events seen at those very long lag times

**Figure 22:**
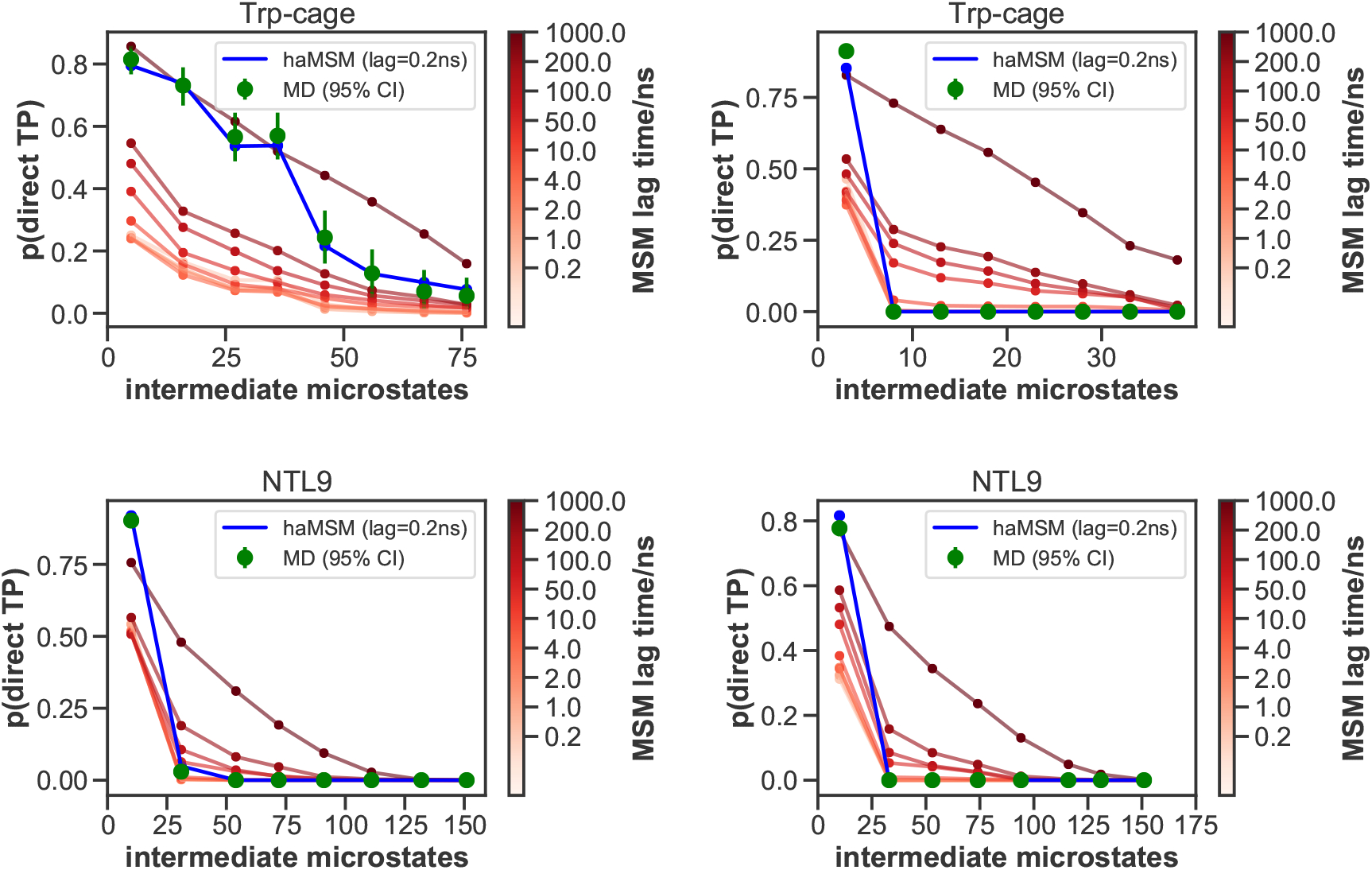
Comparison of top models selected at scoring lag times 100 ns (left side) and 10 ns (right side), for Trp-cage and NTL9: simple mechanism comparison of MD, MSMs, and haMSMs using the fraction of direct folding pathways. For the given number of intermediate microstates, ensembles of discretized transition trajectories were analyzed to determine the fraction which directly ‘hopped over’ the intermediate region based on either the MD discretization time Δ*t*, also used for haMSM modeling, or else the indicated MSM lag time. A greater number of intermediate microstates indicates relatively smaller macrostates and accounts for the monotonic decrease of direct transitions. Two haMSMs data points are missing for NTL9 (10 ns) due to numerical problems.

**Figure 23:**
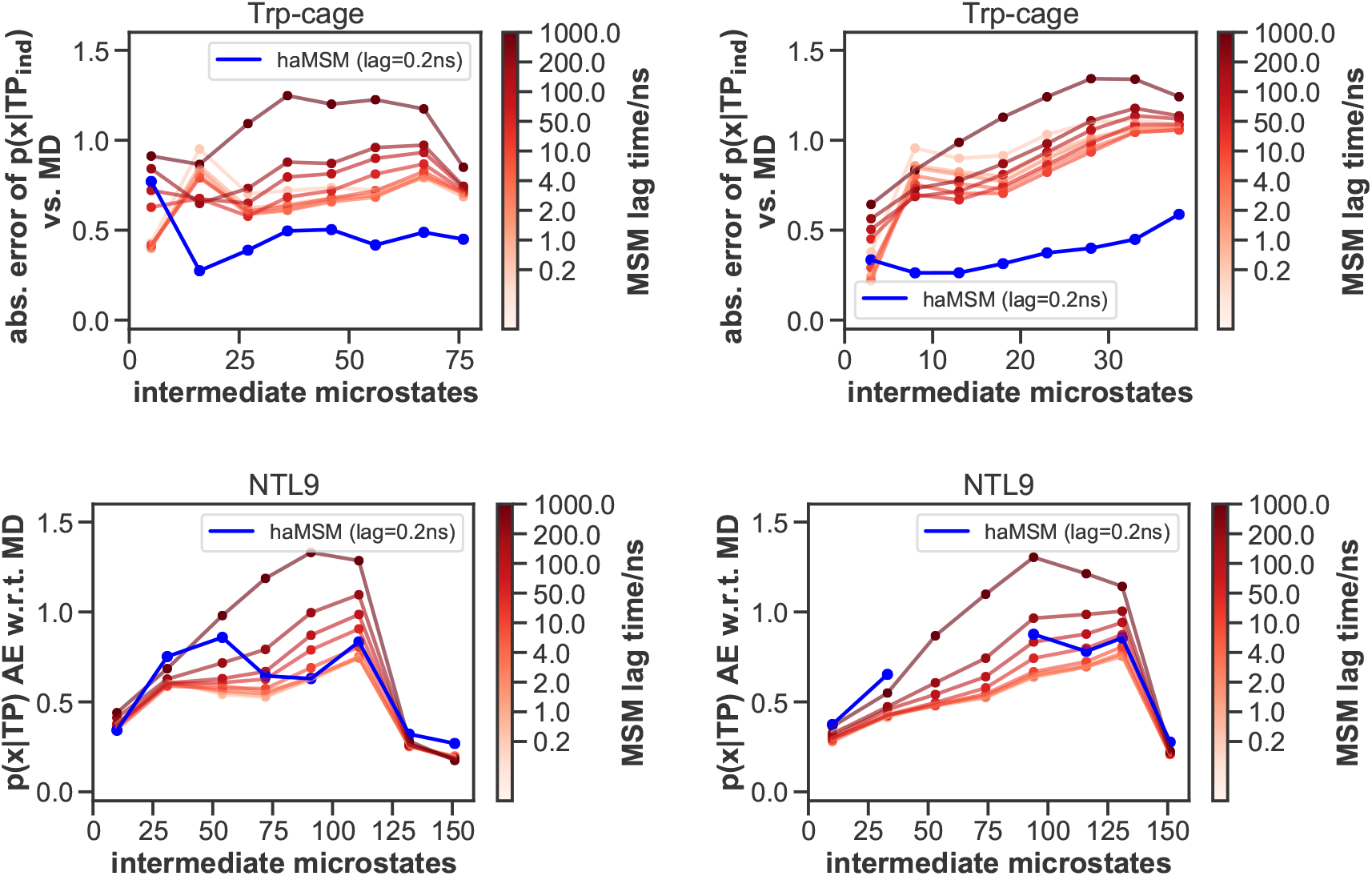
Comparison of top models selected at scoring lag times 100 ns (left side) and 10 ns (right side), for Trp-cage and NTL9: mechanism comparison of MSMs and haMSMs to MD using the configurational distributions of transition path ensemble. Each panel plots the summed *absolute error, as compared to MD,* for intermediate microstate probabilities calculated for the transition path ensembles, i.e., *p*(*x|TP*_ind_), for the given number of intermediate microstates. Importantly, the “ind” subscript indicates that direct pathways analyzed in Figure 6 were excluded from the ensembles prior to computation of errors; had they been included, the MSM errors would be substantially larger. Two haMSMs data points are missing for NTL9 (10 ns) due to numerical problems.

### Defining macrostates based on kinetic clustering

As an alternative coarse-graining procedure, we use a hierarchical (or progressive) clustering based on a cutoff *t_cut_*. If the round-trip time (*t_ij_*) between any two states is less that *t_cut_* then we merge the states. The procedure is as follows:

1. Compute MFPT matrix *M* and add it to *M ^T^* to obtain the round-trip times {*t_ij_*}
2. While min({*t_ij_*}) *< t_cut_*:

- Merge the corresponding states
- Recompute {*t_ij_*} (step 1) for merged states
3. Increase *t_cut_* until clustering results in only one macrostate. Plot the highest {*t_ij_*} vs. *t_cut_*, identify the longest plateau, and take macrostates at *t_cut_* in the middle of the plateau. The following *t_cut_* values were identified: 436 ns for chignolin, 277.6 ns for Trp-cage, 449.2 ns for villin, and 817.6 ns for NTL9.

### Issues with DESRES NTL9 dataset Thermodynamics and kinetics of trajectory NTL9-2 are inconsistent with the other trajectories

While the folded content of the other three NTL9 trajectories is ∼90%, in agreement with the MSM results, trajectory NTL9-2 has only ∼50% folded frames. This can be clearly seen in the RMSD plots of the ‘Individual proteins’ section of the SI of^39^ (NTL9-2 is the 3rd trajectory). The kinetics are also much faster: using macrostates defined from an MSM computed including NTL9-2, this trajectory has 185 folded-unfolded transitions, compared to just 7 in trajectory NTL9-3 of similar length. This is especially pronounced at the beginning of the trajectory—removal of the first 80 microseconds (mean of the folding and unfolding MFPTs in our converged MSM) leaves only 18 transitions left, though still many more than just 6 in the same length of NTL9-3. We removed trajectory NTL9-2 from the dataset analyzed in this paper.

### Topology of trajectory NTL9-1 is different from the other trajectories

The dataset is provided with one NTL9.pdb topology file, as well as .mae files for each trajectory, to use for loading the topology-less .dcd trajectory files. However, using the NTL9.pdb file with the NTL9-1 trajectory provides clearly nonsensical results, e.g. by visual inspection in PyMOL^83^ or lack of MSM convergence if using features defined from this file. Inspection of the .mae files reveals the arrangement of the hydrogen atoms in the topologies are different between NTL9-1 (within each residue, except for the first residue, some of whose hydrogens are at the end of the file) and the other trajectories (at the end of file). Importantly, conversion of the .mae file for NTL9-1 to a .pdb file with PyMOL^83^ or Maestro^88^ FAILS to preserve the original order of hydrogens, while VMD^89^ preserves it and was used by us for the conversion. The results were verified by manual inspection in PyMOL.^83^

## Notes

### Competing Interest Statement

The authors have declared no competing interest.

